# Noncovarying storage effect: balancing and positive directional selection on mutant alleles that amplify random fitness and demographic fluctuations

**DOI:** 10.1101/2024.10.10.617629

**Authors:** Yuseob Kim

**Affiliations:** Division of EcoScience, Ewha Womans University, Seoul, Korea 03760

**Keywords:** fluctuation, storage effect, population size, positive feedback, eco-evolutionary

## Abstract

Temporally variable environments in natural populations generate fluctuations in not only the population size but also the fitness effects of mutant alleles. The theory of storage effect, a species/allelic diversity-promoting mechanism discovered in ecology, predicts that rare mutants with fluctuating fitness can be positively selected and then maintained in balanced polymorphism if the population is subdivided into two parts that are respectively exposed to and protected from fluctuating environment. Recent study found that, under pre-exisiting oscillation in population size, the mutant is positively selected to fixation if its fitness change correlates with the rate of population growth, which further amplifies population size oscillation. To further understand these eco-evolutionary dynamics and elucidate their generality in natural populations, this study built more realistic models that assume randomly, not cyclically, fluctuating selection and common demographic features, including heterogeneous ecological patches or an age-structured population. Mathematical analysis revealed that this novel evolutionary force is generated when the size of the subpopulation subject to weak selection does not covary with that of the other subpopulation. Simulations showed that this ‘noncovarying storage effect’ is robust to various perturbations in the model. Multi-locus simulations revealed that oscillatory polymorphism at many loci can be simultaneously maintained and that positive feedback between demography and selection can accelerate the sequential fixations of fluctuation-amplifying mutations and thus lead to a drastic amplification of population size fluctuation. These results suggest that the noncovarying storage effect is a potentially prevalent evolutionary force in nature for maintaining genetic variation and causing large demographic fluctuations.

**ARTICLE SUMMARY:** Natural populations experience rapid environmental changes that cause fluctuations in the fitness of genetic variants and population size. This study investigates how evolutionary changes due to fitness differences interact with concurrent demographic changes. The theory of the storage effect, which is a mechanism for promoting genetic diversity, was extended to add random population size fluctuation with heterogeneous environmental patches or in a population divided into different age classes. A novel evolutionary force that either increases genetic variation or drives evolution toward further amplification of population size fluctuation, an unconventional contributor to large demographic fluctuations in nature, was identified and analyzed.

## INTRODUCTION

Natural populations experience temporal changes in abiotic and biotic environments. A randomly variable environment is expected to produce fluctuations in the fitness effects of non-neutral or phenotype-changing alleles segregating in the population. If the magnitude of fluctuation in the relative fitness is large, evolutionary changes may occur rapidly at the pace of short-term ecological/demographic fluctuations. Recent developments in evolutionary biology have revealed that such rapid evolutionary changes are widespread (Bell 2010; Thompson 2013; Messer *et al*. 2016; Lynch *et al*. 2024). Temporally variable environments should also generate fluctuations in various aspects of demography, most importantly the number of reproducing individuals (i.e., population size). One may then ask whether mutual interaction between demographic and selective fluctuations creates qualitatively novel eco-evolutionary dynamics that cannot arise if one of the demographic or population genetic variables is held constant.

Considering that a fluctuation in population size reflects a fluctuation in the reproductive success of individuals and that long-term reproductive success is maximized if its variance in the offspring number over time is minimized (Slatkin 1974; Gillespie 1977; Tuljapurkar 1982), one may predict that a population will evolve to reduce size fluctuation as much as possible (Pfister 1998). For example, a variant increasing the egg size rather than the egg number at reproductionwhen plentiful resources are available may be positively selected as the offspring can survive better during an unfavorable period. Then, when a large fluctuation in population density, such as mast seeding or insect outbreak, does occur despite the advantage of a small temporal variance in offspring number, this may be seen as a non-adaptive response of the population to a heavy fluctuation of the biotic and/or abiotic environment (Myers 1988; Dwyer *et al*. 2004; White 2008; Bogdziewicz *et al*. 2020). However, recent studies suggested that a species may evolve to undergo a larger fluctuation in its population size. Liu *et al*. (2019) found that a reproductive strategy increasing the temporal variance of offspring number can be advantageous in populations reproducing in overlapping generations under various patterns of environmental fluctuations. Kim (2023) showed that a mutation amplifying the cyclic fluctuation of population size can be positively selected if this population is partially protected from the cyclical selection (i.e. fitness fluctuation occurs only in a subset of the population).

Partial protection from fluctuating selection commonly occurs in many species. In spatially structured species the amplitude of fluctuation in selective pressure is likely to be heterogeneous over the geographic range. In plants and animals that have their life cycles divided into several stages and reproduce in overlapping generations, selective pressure is unlikely to act uniformly over individuals of all stages or ages. For example, if a trait under fluctuating selection is expressed in the insect larval stage, alleles affecting this trait currently carried by adults are not visible to the selection. Similarly, alleles affecting traits that appear after germination remain neutral in the seed bank of a plant species. Such a subset of the population protected from fluctuating selection may be called a “refuge.” The presence of a refuge of various kinds was found to generate the advantage of rare alleles, a form of balancing selection commonly referred to as the storage effect (Ellner & Hairston 1994; Reinhold 2000; Gulisija & Kim 2015; Gulisija *et al*. 2016; Bertram & Masel 2019; Yamamichi *et al*. 2023), as the refuge acts to temporarily store the variants that otherwise will be removed from the population during unfavorable periods. The storage effect was first theorized in community ecology, shown to arise not only with partial protection from fluctuation provided by a refuge but with general covariance between environment and competition, and is now one of the most important ecological theories for explaining the stable coexistence of species (Chesson & Warner 1981; Chesson 1994; Barabás *et al*. 2018). Imported into population genetics, it was recently proposed as one of a few mechanisms that may produce the genome-wide occurrence of oscillatory polymorphism, as observed in North American *Drosophila melanogaster* and other species, which appear to be maintained under fluctuating environments (Bergland et al. 2014; Park and Kim 2019; Johnson et al. 2023).

However, Kim (2023) showed that if the oscillation of population size is added to the model of storage effect a variant whose fitness also oscillates in the same directions can be either maintained in balanced polymorphism or positively selected to reach fixation. Mathematical analysis showed that whether balancing or directional selection occurs on this fitness-amplifying allele depends on the relative strengths of fitness versus population size fluctuations. Previous studies on balancing selection by the storage effect in a constant-sized population required the coexisting variants to have identical geometric mean fitness. However, when the population size was allowed to oscillate, it led to negative frequency-dependent selection on mutants with even smaller geometric mean fitness than the wild-type allele. It was also shown that positive directional selection on such variants, which cannot arise in the previous models of the storage effect, occurs if demographic fluctuation is relatively stronger than fitness fluctuation. Furthermore, Kim (2023) found that, under a weak regulation of population density under which the population size can over- or undershoot the carrying capacity, the positive selection leads to the amplification of oscillation in mean absolute fitness, thus population size. In summary, the addition of population size fluctuation, either externally imposed by ecology or due to the segregation of fitness-amplifying mutants, in the model was found to generate novel ecoevolutionary dynamics not predicted by the conventional model of the storage effect.

The generality of novel eco-evolutionary dynamics discovered in Kim (2023) may however be questioned since the major result was obtained from the simulation of cyclically (seasonally) and symmetrically changing environments under which the population size and fitness oscillations are fully correlated. Although a mathematical analysis found general conditions for determining which evolutionary outcome (the loss, balanced polymorphism, or fixation of the fitness-amplifying allele) is expected, it was based on the model of only two alternating seasons with fixed sizes. Therefore further investigation using more realistic models is needed to obtain better theoretical understanding and to find out whether the novel eco-evolutionary dynamics can arise in a substantially wider range of biological settings.

This study broadens the scope of investigation in the following directions. First, mathematical models assuming random, rather than deterministic and symmetrically seasonal, fluctuations in the environment were built and analyzed. A model with seasonal fluctuation is suitable for cases in which one cycle of environmental changes spans multiple generations. However, for many species, one generation is not shorter than a seasonal cycle. Random fluctuation of carrying capacities and the direction/strength of selection over generations, with a varying degree of correlation between these fluctuations, is probably relevant to a wider range of species including plants that exhibit mast seeding. Second, new models of population structures and inter-deme differences in selection intensity were investigated. Whereas Kim (2023) focused on a population with a seed bank (Turelli *et al*. 2001; Gremer and Venable 2014) in which the mutant is completely protected from selection in the refuge, this study uses a model (the two-patch or TP model) that assumes two demes or ecological patches, both reproducing and undergoing changes in the carrying capacity, under general heterogeneity in selection intensity. This TP model was also analyzed extensively by simulation to test its robustness against perturbations. Another model (the larval-subadult-adult or LSA model) assuming a population with an age structure, in which the life-cycle is divided into the larval (juvenile), subadult, and adult stages, was also explored by simulation. Third, for both TP and LSA models, multi-locus stochastic simulations were performed. Simulations showed that mutant alleles causing wider fluctuation in offspring number can reach fixation at multiple loci in a wide range of conditions. These fixations were found to occur at faster rates with increasing numbers of loci, as fluctuating population size and fluctuating selection reinforced each other in positive feedback.

As will be clarified in the mathematical analysis, a key ingredient for this novel eco-evolutionary dynamics is the presence of a refuge whose size (or carrying capacity) remains relatively invariant or does not covary with the size of the rest of the population. One may therefore call this evolutionary mechanism the “noncovarying storage effect”. The result suggests that the noncovarying storage effect arising under the general condition of heterogeneous fluctuating selection in structured populations likely contributes to maintaining non-neutral genetic variation and to increasing demographic variability. Analytical and simulation results for the TP model will be first presented, followed by simulation results for the LSA model.

### Two-patch (TP) model

Consider a haploid population that reproduces in discrete generations. The outline of the life cycle is illustrated in Fig. 1A. The population is divided into two subpopulations or patches that are subject to different levels and/or phases of fluctuating selection (thus two-patch or TP model). For convenience, these subpopulations will be called the field and the refuge following the terminology of Kim (2023). The latter refers to a subpopulation where selection on a mutant allele is weaker. It is assumed that the population is initially fixed for the ancestral *A*_1_ allele at a locus. Then, a new mutant allele *A*_2_, subject to fluctuating selection arises. The fitness of an individual carrying *A*_2_ relative to *A*_1_ is given by 1 + *S*_*t*_ in the field and 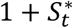in the refuge, where 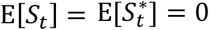, Var[*S*_*t*_] > 0, and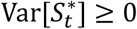. Notably, while the arithmetic mean of the fitness of *A*_2_ over time is 1, its geometric mean is less than 1, in both subpopulations. Natural selection is thus expected to eliminate such an allele from a simple panmictic (unstructured) population.

**Figure 1.**
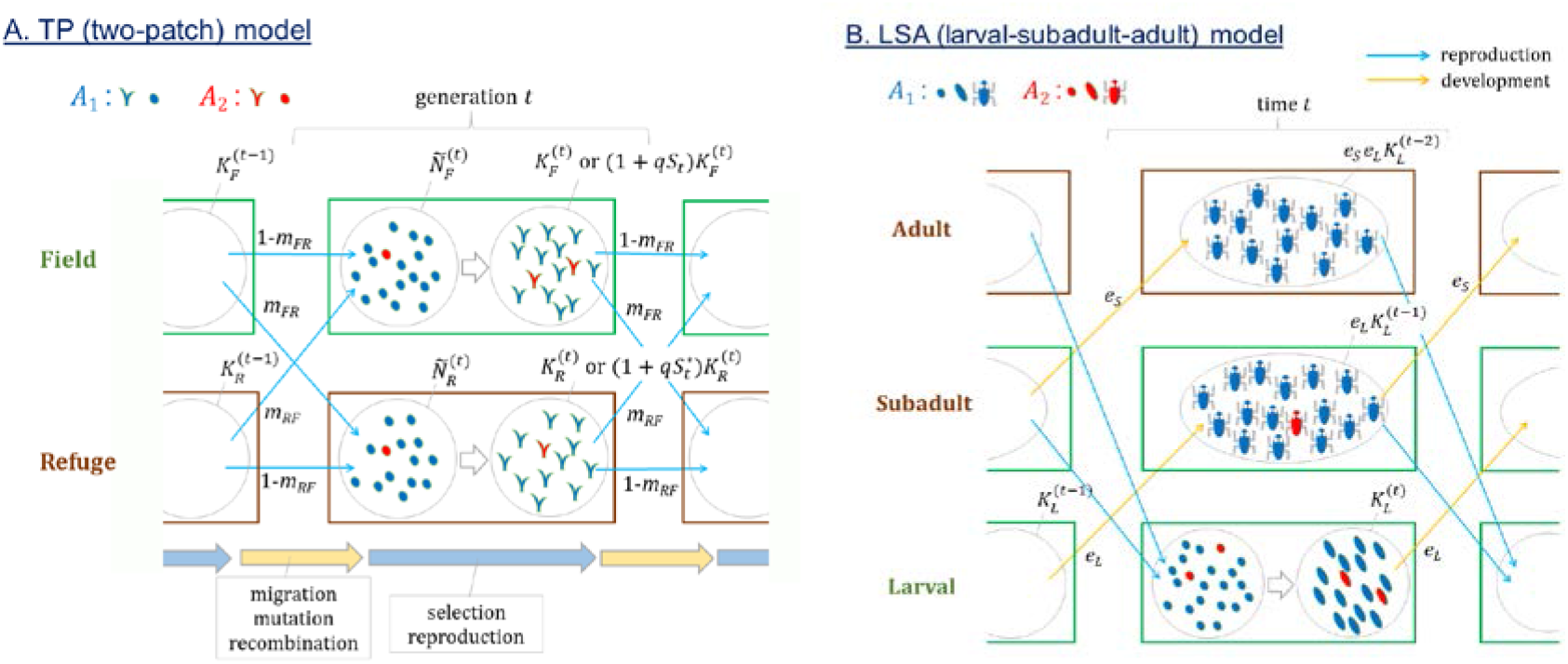
Schematic diagrams of population models. **A**. Two−patch (TP) model assumes a population of haploid individuals occupying a field, where the mutant allele *A*_2_ has selective advantage 1 + *S*_*t*_ in generation *t*, and a refuge, where the advantage is 1 + *S*^*^_*t*_. Each generation starts with young individuals that either migrated across or stayed in subpopulations after being reproduced in the previous generation. They also undergo mutation and recombination before selection starts. Then, they undergo selection such that the total number of reproducing individual is constrained to the carrying capacity 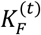for the field and 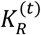for the refuge assuming soft selection. In case of hard selection the carrying capcities are effectively 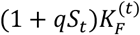and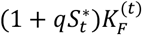. **B**. In the larval−subadult−adult (LSA) model, at each time step haploid individuals move from larval to subadult stage with probability *e*_*L*_ and from subadult to adult stage with probability *e*_*S*_. Reproduction, followed immediately by mutation and reproduction, occurs only in the subadult and adult stages. A new larval subpopulation at time *t* starts with individuals produced at time *t*−1. At the end of time interval *t*,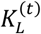 individuals remain in the larval stage before moving to the next stage.

A fluctuating environment causes fluctuations in the carrying capacities of both the field and refuge, which are given by 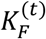and 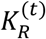respectively for generation *t*. The carrying capacity is defined as the expected number of (reproductive) individuals at the end of each generation. It is assumed that the carrying capacities of both subpopulations depend on common environmental factors that fluctuate randomly over time but that subpopulations may differ in the strengths of the dependence. The sizes of subpopulations, 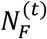and 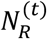, after the reproduction step in generation *t*, may or many not match the corresponding carrying capacities depending on the strength of population density regulation (see below). Before a new generation starts, each individual in the field migrates to the refuge or remains in the field with probability *m*_*FR*_ and 1 - *m*_*FR*_. Similarly, each individual in the refuge either migrates or not with probability *m*_*RF*_ and 1 - *m*_*RF*_.

Next, the size of the field (refuge) changes from 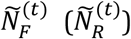 at the beginning of generation *t* to 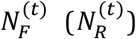 at the end, where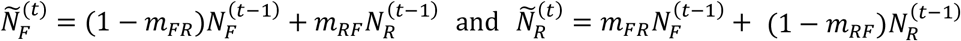.

Let *n*_1F_ and *n*_2F_ be the numbers of individuals in the field carrying *A*_1_ and *A*_2_ alleles at the beginning (before the reproduction step) of generation *t* (thus 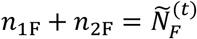). In the reproduction step, two different assumptions can be made regarding the effect of the mean fitness of individuals on the total population size. When the allele with higher or lower fitness increases in frequency, the total population size may not change due to an ecological constraint. In other words, the absolute fitness of the allele is frequency-dependent while its relative fitness remains constant. Conversely the mutant allele may increase or decrease the absolute reproductive output of the carriers regardless of its frequency such that the population size can increase over or decrease below the carrying capacity, which is now interpreted as the expected population size assuming that all individuals carry the ancestral (*A*_1_) allele. For convenience, these two modes of natural selection will be simply referred to as soft and hard selection, respectively, following the terminology in Wallace (1975). First, under soft selection, very strong density regulation occurs so that the final size of the field is constrained to 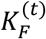. The number of progeny born to each parent is given by 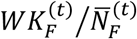, where the fitness *W* is 1 for *A*_1_ and 1 + *S* for *A*_2_, and 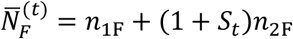.

Second, under hard selection, fitness-changing mutations lead to changes in not only allele frequencies but also in the population size. To this effect, the number of progeny per parent is now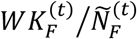. Namely, when *q* = *n*_2F_/(*n*_1F_+ *n*_2F_) is the relative frequency of *A*_2_ at the beginning of a generation, the size of the field is expected to reach 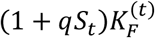, which becomes the effective carrying capacity of the field under hard selection. Similarly, reproduction in the refuge occurs either with soft or hard selection, where subscript *F* in the above expressions is replaced by *R* and *W* is 1 for *A*1 and 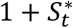 for *A*2. The model so far described changes in the number of individuals without genetic drift. When a finite population size is assumed, as in simulation below, each parent in both subpopulations is modeled to produce a Poisson number of offspring with the mean specified above, which approximates genetic drift in the Wright-Fisher model.

Finally, how the environment fluctuates over time and how demographic and selective parameters —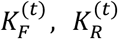, *S*_*t*_, and 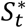— respond need to be specified. The environmental condition is assumed to fluctuate randomly in each generation, therefore with no temporal autocorrelation.

More specifically, fluctuations occur such that 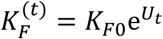 and 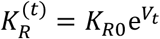 where *U*_*t*_ and *V*_*t*_ are drawn each generation from a joint normal distribution satisfying E[*U*_*t*_] = E[*V*_*t*_] = 0. The oscillation of carrying capacities is therefore symmetrical in the log scale, which is generally predicted in demographic models and observed in nature (Tuljapurkar & Orzack 1980; Myers 1988). It is assumed that *S*_*t*_, 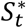, *U*_*t*_, and *V*_*t*_ may be correlated to each other as they respond to the fluctuation of common environmental variables. However, the correlation among them satisfies that Cov[*S*_*t*_, *U*_*t*_ − *V*_*t*_] is greater than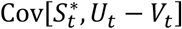, which gives the mathematical definition of the field and refuge.

### Analytic results: the fate of the *A*_2_ allele

This section aims to predict the fate of the rare *A*_2_ allele — whether it will be lost, fixed, or increased to intermediate frequencies and maintained polymorphic in the population — as a function of the demographic and fitness parameters in the TP model. For convenience in the derivation, genetic drift is ignored and a fixed scheme of migration, *m*_*FR*_ = 1 − *m*_*RF*_ = *r*, that sends a constant fraction *r* of the population to the refuge regardless of the current location is assumed. (Below the dynamics will be examined in simulation without this restriction in the migration rate.) First, soft selection, under which 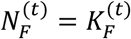 and 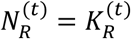, is assumed. Let *n*_1_ and *n*_2_ be the total numbers of individuals (counted after reproduction/selection and before migration) carrying *A*_1_ and *A*_2_ in generation *t* – 1 and *n’*_1_ and *n’*_2_ be their expected numbers in generation *t*. Then, with *n*_1_ ≫ *n*_2_,

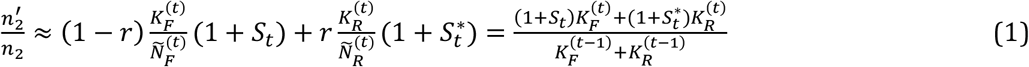

(Appendix A). This equation takes into account that the change in the absolute number of *A*_2_ is determined first by demography, as the size of field changes from 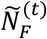 to 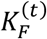 and that of refuge from 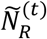 to 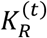, and also by fitness 1 + *S*_*t*_ and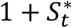. Whether the copy number of rare allele *A*_2_ is expected to increase or decrease is determined by the expectation of 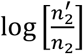over demographic and fitness fluctuations. It is shown in Appendix A that the condition for the *A*_2_ allele being positively selected (i.e. successfully invading the population initially fixed for *A*_1_), 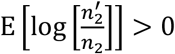, is simplified to

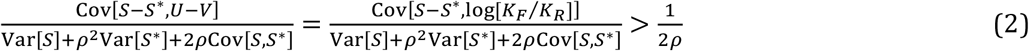

where *ρ* = *K*_*R*0_/*K*_*F*0_ (note that 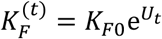 and 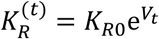 represents the relative size of the refuge and super-/subscripts for time *t* in variables were omitted.

If it is assumed that selection on *A*_2_ in the refuge is weaker than that in the field by a ratio δ, namely 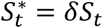, the above inequality is simplified to

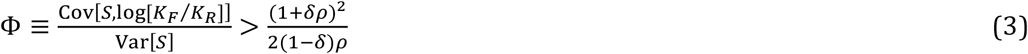

Therefore, the rare *A*_2_ allele is positively selected if the covariance between its fitness and the log-ratio of the field to refuge size is large relative to the amplitude of the fitness fluctuation. This requires that the demographic fluctuation in the population be sufficiently large, as already shown in Kim (2023), and that, to allow the substantial fluctuation of 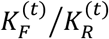, the carrying capacity of the refuge should not covary strongly with that of the field. Since this requirement for the refuge is essential and distinguishes the current from the previous models of storage effect, the evolutionary force driving this positive selection may be called the noncovarying storage effect (see Discussion for examples of noncovarying and covarying storages). The conditions (2) and (3) are easier to meet as δ becomes small well below 1, namely when the fitness fluctuation of *A*_2_ becomes either smaller in the refuge compared to the field or negatively correlated to *S*_*t*_.

In a similar manner, the condition for the *A*_1_ allele to invade a population initially fixed for *A*_2_ under soft selection becomes

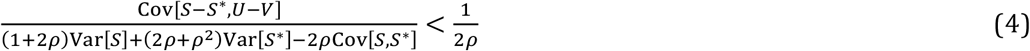

(Appendix A) or

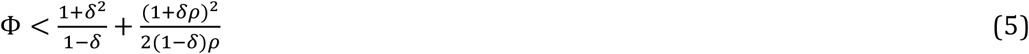

If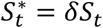.

In a simpler case with δ = 0, from inequalities (3) and (5), it is predicted that the *A*_1_ and *A*_2_ alleles coexist in the population if 1/(2*ρ*) < Φ < 1 + 1/(2*ρ*) under soft selection. With Φ < 1/(2*ρ*), the *A*_2_ allele is eliminated from the population, as the fitness fluctuation that reduces its geometric mean in the field is more important than population size fluctuation. With Φ > 1 + 1/(2*ρ*), the *A*_2_ allele is predicted to reach fixation in the population.

Using the same approach, the fate of the *A*_2_ allele under hard selection can be analyzed. Note that the carrying capacities of the field and refuge are modified to 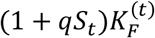and 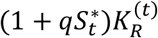 with hard selection where *q* is the frequency of *A*_2_. The frequency change of *A*_2_ when rare is therefore effectively identical to that in soft selection. Therefore, the condition for *A*_2_ invading the population fixed for *A*_1_ is again given by inequalities (2) and (3). However, the condition for the fixation of *A*_2_ is drastically different from that under soft selection. It is shown in Appendix A that the condition for 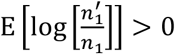 for rare *A*_1_ is exactly the condition for 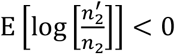for rare *A*_2_. Therefore, with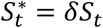, the condition for the *A*_1_ allele to invade the population fixed for *A*_2_ is approximately Φ < (1 + δ*ρ*)^2^/(2(1 − δ)*ρ*). This means that the fixation of *A*_2_ is ensured if Φ > (1 + δ*ρ*)^2^/(2(1 − δ)*ρ*); once positively selected after its appearance by mutation, the *A*_2_ allele is expected to increase in frequency all the way to fixation. This leads to the amplification of fluctuation in the size of the field, from 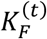to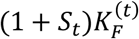. This result is consistent with the finding in Kim (2023) that, under weak density regulation of the population size, an allele with a wider oscillation in absolute fitness reaches fixation in the presence of refuge.

### One-locus simulation

To validate the above analysis and explore further eco-evolutionary dynamics in the TP model, the one-locus stochastic simulation was performed. The demographic and fitness fluctuations were modeled by *U*_*t*_ = *a*_*U*_*Z*_1_ + *b*_*U*_*Z*_2_, *V*_*t*_ = *a*_*V*_ *Z*_1_ + *b*_*V*_*Z*_3_, *S*_*t*_ = *a*_*S*_*Z*_1_ + *b*_*S*_*Z*_4_, and 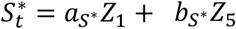 where *a*_*X*_ (*b*_*X*_) is a constant for determining the magnitude of change in variable *X* due to a common environmental factor (other factors) and the *Z*_*i*_s are standard normal variates that were drawn independently each generation. Therefore, *Z*_1_ represents the state of the environment affecting both population sizes and fitness.

Simulations were first performed for a scenario in which *A*_2_ allele is completely protected from selection in the refuge 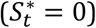 and migration follows a simple scheme (*m*_*FR*_ = *m*_*RF*_ = 0.5). As suggested by the analysis above, the major parameter of eco-evolutionary dynamics in this case is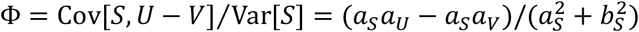. Simulation results were obtained for various values of Φ under soft (Fig. 2A, B) and hard (Fig. 2C, D) selection. Initially the numbers of *A*_1_ and *A*_2_ individuals changed deterministically once the demographic and fitness parameters (*U, V*, and *S*) for a given generation were sampled from their joint distribution, effectively assuming that the sizes of the field and refuge were finite and fluctuating but very large so that genetic drift can be ignored (Fig. 2A). Various values of *a*_*S*_ and *b*_*S*_ were chosen so that Φ ranged from 0.3 to 2, while ρ was set to 1. Starting from 0.5, the frequency of the *A*_2_ allele at the 5,000th generation was recorded for 1000 replicates per parameter set. As predicted, with Φ ≤ 0.5, the frequency of *A*_2_ approached 0, thus confirming negative selection, and with Φ ≥ 1.5 the frequency approached 1, thus confirming positive directional selection. The distribution of the allele frequency was approximately uniform between 0 and 1 when Φ = 1. Next, genetic drift and bidirectional mutation between two alleles were added to the above simulations (Fig. 2B). This weakened the strengths of all modes of selection. With Φ = 1, more simulation runs ended with allele frequencies close to 0 and 1. However, the proportion of runs ending with intermediate allele frequencies was still larger than that observed when the alleles were set as neutral (dashed curve in Fig. 2B).

**Figure 2.**
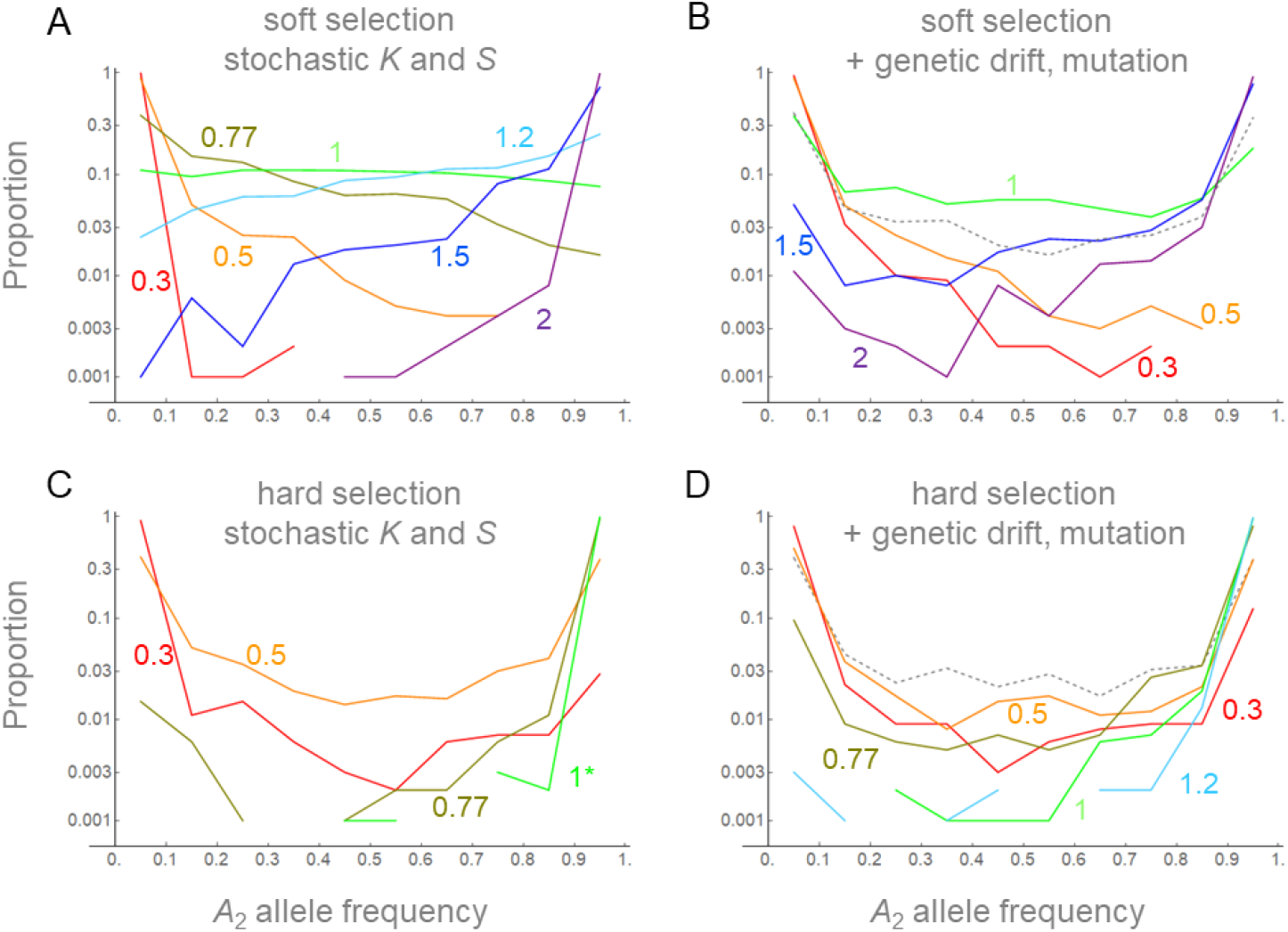
Distribution of *A*_2_ allele frequencies (*p*) at the 5,000^th^ generation in the stochastic simulations of soft (A, B) and hard (C, D) selection in the TP model. The initial frequency of *A*_2_ is 0.5 in both field and refuge. Reproduction occurred while 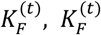and *S*_*t*_ varied stochastically without (A, C) or with (B, D) genetic drift and bidirectional mutation (μ = 5×10^−5^). Proportions of simulation replicates (out of 1000) with the frequencies in the interval 0 ≤ *p* < 0.1, 0.1 ≤ *p* < 0.2, …, 0.8 ≤ *p* < 0.9, and 0.9 ≤ *p* ≤ 1 were plotted and connected by lines. Fluctuations of population sizes are given by *K*_*R*0_ = *K*_*F*0_ = 1000 (*ρ* = 1), *a*_*U*_ = 0.3, *b*_*U*_ = 0.1, *a*_*V*_ = 0.05, *b*_*V*_ = 0.1. Parameters of fluctuation were chosen to yield Φ = 0.3, 0.5, 0.77, 1, 1.2, 1.5, and 2: (*a*_*S*_, *b*_*S*_) = (0.07, 0.23), (0.1, 0.2), (0.14, 0.16), (0.2, 0.1), (0.2, 0.04), (0.15, 0.05), and (0.1, 0.05). In the case of hard selection with Φ = 1 and no genetic drift (panel C), simulation ran only upto 2,500 generations: longer simulations or larger Φ resulted in all replicates with final frequencies > 0.9. The results connected by dashed lines in B and D represent the cases of neutral evolution (*a*_*S*_ = *b*_*S*_ =0).

Therefore, an evolutionary force to maintain polymorphism beyond the level under the neutrality, namely balancing selection, was confirmed.

Similarly, the fate of the *A*_2_ allele under hard selection was examined. Stochastic simulation confirmed the approximation above predicting that *A*_2_ is positively selected towards fixation if 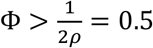, although genetic drift and mutation weakened the trend (Fig 2C,D). Positive selection on both rare and common *A*_2_ under the same condition means that this allele may not remain polymorphic for long in the population. Indeed, across all values of Φ there were fewer simulation runs ending with intermediate frequencies of *A*_2_ compared to the simulation of neutral alleles subject to genetic drift and bidirectional mutations only (dashed curve in Fig. 2D). Therefore, balancing selection that was observed with soft selection did not arise with hard selection.

Next, the cases in which *A*_2_ allele is not completely neutral in the refuge was examined. A simple model setting 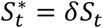was used. This time, the *A*_2_ allele is initially absent in the population but introduced with recurrent mutation. For a given parameter set simulation was run over at least 10^5^ generations and the mean 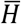of expected heterozygosity (2*q*(1 − *q*), where *q* is the relative frequency of *A*_2_) and the proportion of time when 0 ≤ *q* < 0.1, *P*_01_, and 0.9 < *q* ≤ 1, *P*_90_, during simulation were recorded. Simulation with 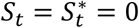 but under the same demographic structure yielded 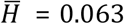(average over 6×10_5_ generations). Then, as balancing selection is defined to be a force yielding genetic variation above the neutral level, a result was classified as balanced polymorphism if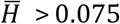. If 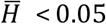and *P*_01_ > *P*_90_, it suggests that negative selection against *A*_2_ was the dominant evolutionary force. Such results were therefore classified as the loss of *A*_2_, since recurrent losses of *A*_2_ should be the major events in the simulation (confirmed by visual inspection of allele frequency trajectories; not shown). Conversely, results yielding 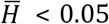and *P*_90_ > 0.7, which indicates that positive directional selection on *A*_2_ was the dominant force, were classified as the fixation of *A*_2_. All others were classified as not distinguishable from the neutral evolution. Simulations largely confirmed the analytic approximations for the conditions for the loss, fixation, or balanced polymorphism of *A*_2_ in a population initially fixed for *A*_1_ (e.g., 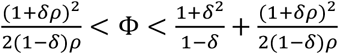 for the balanced polymorphism under soft selection and 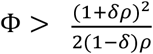 for the fixation under hard selection; Fig. 3). It is shown that increasing δ above 0 makes it harder for balancing and positive directional selection on *A*_2_ to occur; for a large value of δ, the fluctuation (variance) of *S*_t_ needs to be much smaller than its covariance with the fluctuation of 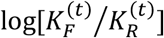 (eq. 3). Small Var[*S*_*t*_] then means that the relative strength of genetic drift affecting the evolutionary dynamics of *A*_2_ is large, explaining the results not distinguisable from the neutrality with large δ and Φ in Figure 3. Other results not distinguishable from neutral evolution were obtained when the predicted effects are near the boundary between balancing and directional selection. Another important result is that negative δ, thus selection in the refuge being in the opposite direction, widens the parameter range for balancing selection.

**Figure 3.**
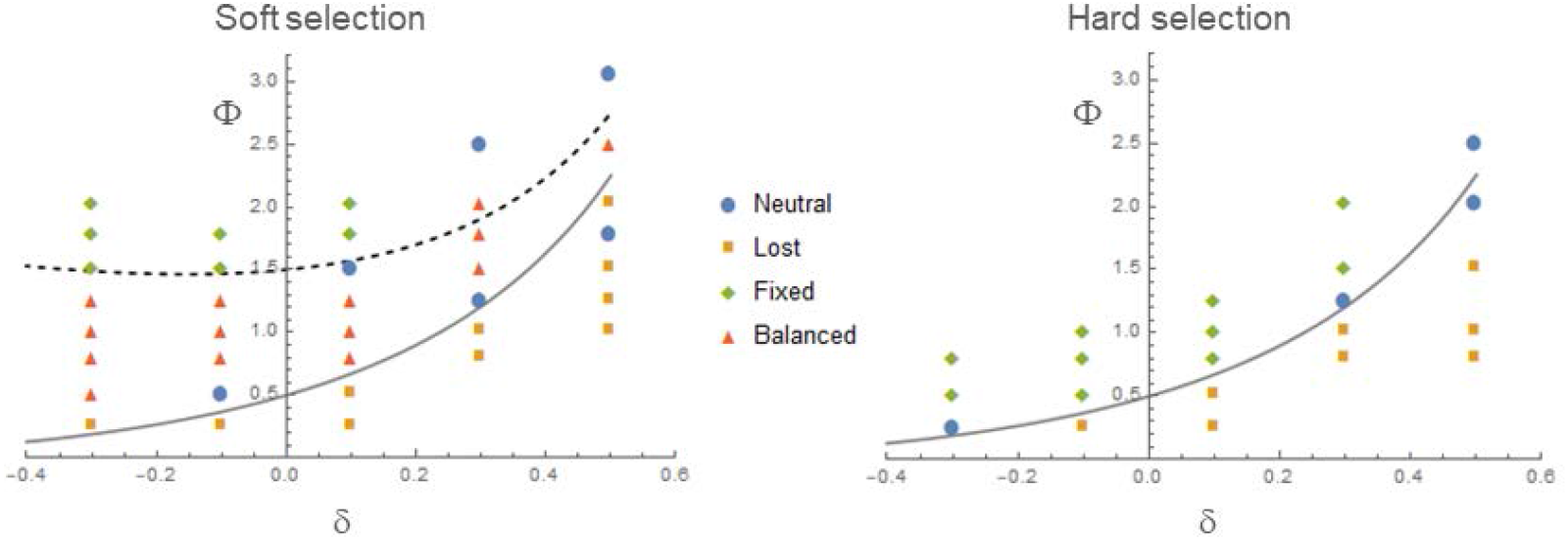
Simulation results for non−zero values of δ. For a given parameter set simulation started with the *A*_1_ allele fixed in the population and ran for 10^5^ generations with recurrent bidirectional mutations and soft (left) or hard (right) selection. Four different outcomes regarding the overall behavior of the *A*_2_ allele (balanced, lost, fixed, neutral) were marked by different symbols and plotted as functions of δ and Φ. The demographic parameters are identical to those in Figure 1. Five different values of δ (−0.3, −0.1, 0.1, 0.3, and 0.3) and ten different values of Φ were chosen: 0.246, 0.5, 0.8, 1, 1.25, 1.5, 1.786, 2.027, 2.5 and 2.94: (*a*_*S*_, *b*_*S*_) = (0.05, 0.22), (0.1, 0.2), (0.2, 0.15), (0.2, 0.1), (0.2, 0), (0.15, 0.05), (0.14, 0), (0.12, 0.02), (0.1, 0) and (0.085, 0). Gray and dashed curves plot 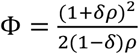and 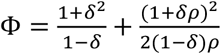with *ρ* = 1.

Finally, the robustness of balancing and positive directional selection above to deviations from the simple migration scheme, *m*_*FR*_ = 1 − *m*_*RF*_ = *r*, was examined. In actual populations, migration rates may vary over time and may not be symmetrical between the field and refuge, at least in a short time scale. New simulations therefore sampled migration rates *m*_FR_ and *m*_RF_ from uniform distributions in ranges [*m*_1_, *m*_2_] and [*m*_3_, *m*_4_], respectively, each generation (Table S1). With ρ = 1 (*K*_*F*0_ = *K*_*R*0_), limiting both *m*_FR_ and *m*_RF_ close to 0.5 yielded the highest level of balanced polymorphism under soft selection. However, allowing random deviation from 0.5 (e.g. [*m*_1_, *m*_2_] = [*m*_3_, *m*_4_] = [0.3, 0.7]) only slightly lowered the level of polymorphism. Smaller migration rates generally disrupted balancing selection. Interestingly, the effect of *m*_FR_ and *m*_RF_ was not symmetric; balancing selection remained effective with *m*_RF_ close to 0.5 and *m*_FR_ less than 0.2, however not with the opposite ratio of rates (compare cases with [[*m*_1_, *m*_2_], [*m*_3_, *m*_4_]] = [[0.1, 0.2], [0.4, 0.5]] and [[0.4, 0.5], [0.1, 0.2]] in Table S1). Similar effects of *m*_FR_ and *m*_RF_ on the directional increase of *A*_2_ allele frequency under hard selection were observed.

In summary, one-locus simulations for the TP model not only confirmed the analytic approximations for the conditions under which the noncovarying storage effect leads to either balanced polymorphism or the fixation of the fluctuation-amplifying allele but also showed that the effect is robust to perturbations in the model such as genetic drift and randomness/asymmetry in the migration rate.

### Multi-locus simulation

To investigate how the eco-evolutionary dynamics generated by the noncovarying storage effect in the TP model play out when there are many such loci in the genome, new sets of simulations were performed in which a haploid individual is modeled to have *L* linked loci, each carrying either the *A*_1_ or *A*_2_ allele described above. To avoid exploring too many parameter dimensions while focusing on the effect of increasing the number of loci, 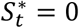and a fixed migration scheme (*m*_*FR*_ = *m*_*RF*_ = 0.5) are assumed. The simulation tracks the numbers of individuals of different haplotypes (see Appendix B). Epistasis is not considered; the relative fitness of haplotype *i* is given by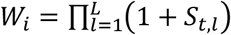, where *S*_*t,l*_ is 0 if the *l*th locus of the haplotype carries *A*_1_ and *a*_*S*_*Z*_1_ + *b*_*S*_*Z*_4_ if *A*_2_. *Z*_1_ is drawn once for all loci while *Z*_4_ is drawn separately for different loci. Individuals randomly pair and undergo recombination by crossing-over such that the probability of recombination between adjacent loci is *c* per generation. The simulation starts with a population fixed for *A*_1_ at all loci. Then *A*_2_ arises by mutations with probability μ/locus/generation.

Two major outcomes of the noncovarying storage effect – balancing selection and positive directional selection – were addressed. First, considering that (1) the negative frequency-dependent selection on the *A*_2_ allele was caused primarily by its fitness correlated to the fluctuation in the carrying capacity (population size) of the field and that (2) this demographic fluctuation should be “felt” at all loci in the genome, one may expect that such balancing selection can occur simultaneously at many loci harboring such variants. To verify if this is true, simulations were run under soft selection and demographic/selective parameters that promoted polymorphism in the single-locus model (Φ = 1 for ρ = 1) with *L* = 1, 4, or 10. Exemplary trajectories of allele frequencies are shown in Fig. S1. Each run lasted at least 10_5_ generations over Which 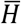, the expected heterozygosity (2*p* (1 − *p*)) averaged over time and loci, and the proportion of time the *A*_2_ alleles spent in frequency between 0.1 and 0.5, *P*_15_, and between 0.5 and 0.9, *P*_59_, were recorded (Table 1). In all cases with fluctuating fitness (*a*_*S*_ > 0), 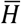was above the level observed in equivalent simulations with neutral alleles (*S*_*t*_ = 0), thus confirming multi-locus balancing selection. For cases of stronger selection (larger Var[*S*_*t*_]), the level of polymorphism per locus declined as *L* increased from 1 to 10. The combined effects of the number of loci and the rate of recombination were not easy to interpret as they depended on the parameters of selection (for sets of fitness/demographic parameters that yield the same Φ = 1, 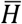 responded differently with varying *L* and *c*). One possible explanation for generally lower variation with 10 loci instead of 1 or 4 is that the simultaneous segregation of alleles at many loci increased the variance of the offspring numbers, thus decreasing the effective population size and weakening the negative frequency-dependent selection that is needed for protected polymorphism. Further exploration of multi-locus polymorphism at a larger scale (in both number of loci and population size) may not be feasible with the current simulation approach based on haplotype frequencies.

**Table 1.** The long-term level of polymorphism in the multi-locus simulation.

| demography | selection mode | $L$ | $c$ | $\bar{H}$ | $P_{15}$ | $P_{59}$ |
| --- | --- | --- | --- | --- | --- | --- |
| $a_U = 0.2,$<br>$a_V = b_U = b_V = 0.1$ | neutral<br>($a_S = b_S = 0$) | 1 | - | 0.0639 | 0.069 | 0.074 |
| | soft selection, $\Phi = 1$<br>($a_S = 0.1, b_S = 0$) | 1 | - | 0.132 | 0.15 | 0.16 |
| | | 4 | $10^{-4}$ | 0.0960 | 0.099 | 0.12 |
| | | 4 | $10^{-3}$ | 0.103 | 0.12 | 0.12 |
|  |  | 4 | 0.01 | 0.0977 | 0.11 | 0.12 |
|  |  | 4 | 0.1 | 0.136 | 0.11 | 0.14 |
| | | 10 | $10^{-4}$ | 0.080 | 0.087 | 0.095 |
| | | 10 | $10^{-3}$ | 0.102 | 0.12 | 0.12 |
|  |  | 10 | 0.01 | 0.109 | 0.15 | 0.10 |
|  |  | 10 | 0.1 | 0.107 | 0.13 | 0.12 |
| $a_U = 0.3, a_V = 0.05,$<br>$b_U = b_V = 0.1$ | neutral<br>($a_S = b_S = 0$) | 1 | - | 0.0633 | 0.081 | 0.074 |
| | soft selection, $\Phi = 1$<br>($a_S = 0.2, b_S = 0.1$) | 1 | - | 0.177 | 0.24 | 0.18 |
| | | 4 | $10^{-4}$ | 0.162 | 0.19 | 0.19 |
| | | 4 | $10^{-3}$ | 0.130 | 0.21 | 0.11 |
|  |  | 4 | 0.01 | 0.168 | 0.22 | 0.18 |
|  |  | 4 | 0.1 | 0.183 | 0.25 | 0.19 |
| | | 10 | $10^{-4}$ | 0.120 | 0.14 | 0.14 |
| | | 10 | $10^{-3}$ | 0.141 | 0.17 | 0.17 |
|  |  | 10 | 0.01 | 0.0934 | 0.076 | 0.14 |
|  |  | 10 | 0.1 | 0.105 | 0.093 | 0.15 |
| $a_U = 0.05,$<br>$a_V = b_U = b_V = 0$ | neutral<br>($a_S = b_S = 0$) | 1 | - | 0.0660 | 0.085 | 0.063 |
| | soft selection, $\Phi = 1$<br>($a_S = 0.05, b_S = 0$) | 1 | - | 0.0816 | 0.11 | 0.081 |
| | | 10 | $10^{-4}$ | 0.0728 | 0.085 | 0.079 |
| | | 10 | $10^{-3}$ | 0.0982 | 0.12 | 0.11 |
|  |  | 10 | 0.01 | 0.0953 | 0.12 | 0.096 |
|  |  | 10 | 0.1 | 0.0952 | 0.13 | 0.092 |

- Simulations were run for $5 \times 10^5$ , $2 \times 10^5$ , and $10^5$ generations with $L = 1, 4$ , and $10$ , respectively.
- Other parameters: $K_{R0} = K_{P0} = 1,000$ ( $\rho = 1$ ), $\mu = 2 \times 10^{-5}$ .

Second, simulation was performed with parameter values under which directional selection for the *A*_2_ allele is expected in the single-locus model (Φ > 1 + 1/(2*ρ*) for soft selection and Φ > 1/(2*ρ*) for hard selection). As expected, substitutions occurred sequentially until *A*_2_ reached fixation at all *L* loci (Fig. 4A and Fig. S2 for example). Times (in generations) when the frequency of *A*_2_ first exceeded 0.9 at each locus were recorded and sorted into *T*_1_ to *T*_*L*_. There is a large difference in substitution dynamics between soft and hard selection. With soft selection, the mean waiting time, 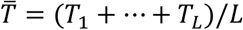, increased moderately as *L* increased from 1 to 8 (Fig. 5A). This may be explained by a reduction in the efficacy of positive selection on *A*_2_ at a given locus when selection at other loci results in genome-wide reduction in the effective population size.

**Figure 4.**
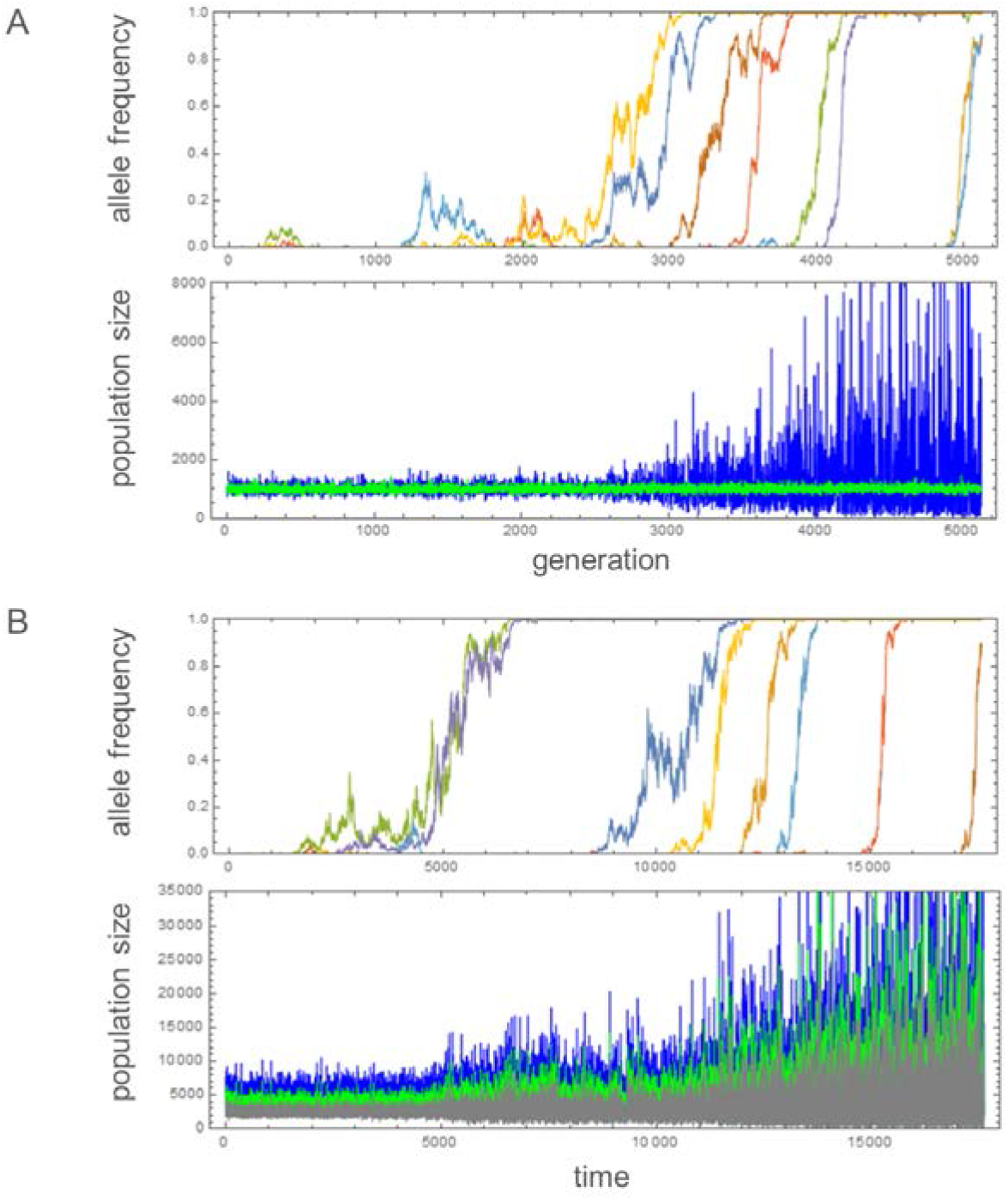
Changes of mutant (*A*_2_) allele frequencies and subpopulation sizes (A. TP model, blue: 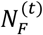, green: 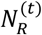; B. LSA model, blue: larval, green: subadult, gray: adult population) in the multi−locus simulations of hard selection. Allele frequency trajectories of 8 loci were plotted in different colors. Parameters: A (TP model): *L* = 8, *c* = 0.02, *K*_*R*0_ = *K*_*F*0_ = 1000, μ = 2×10^−5^, and Φ = 1 (*a*_*S*_ = 0.15, *b*_*S*_ = 0, *a*_*U*_ = 0.15, *a*_*V*_ = 0, *b*_*U*_ =0, *b*_*V*_ = 0.1). B (LSA model): *L* = 8, *c* = 0.02, *K*_*L*0_ = 5000, μ = 2×10^−5^, and Φ’ = 1 (*a*_*S*_ = 0.1, *b*_*S*_ = 0.1, *a*_*L*_ = 0.2, *b*_*L*_= 0.1).

**Figure 5.**
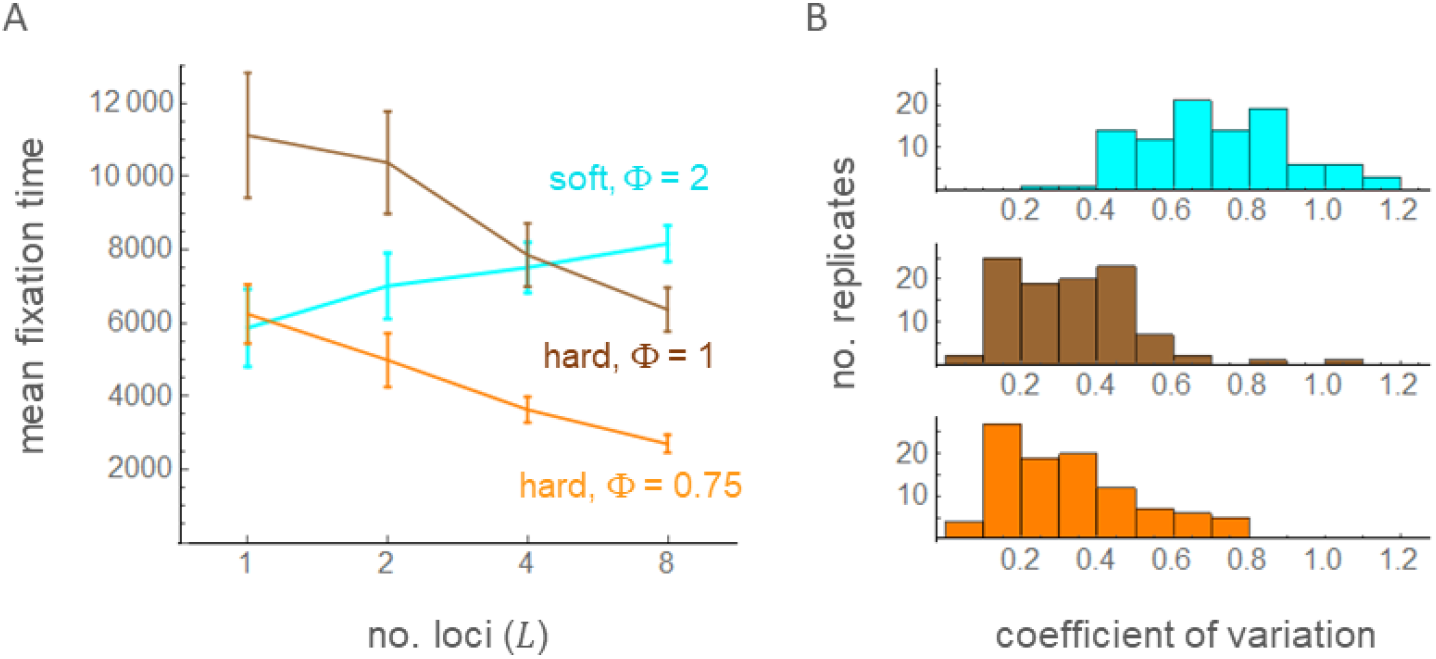
Mean fixation time 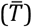 with varying number of loci (A; *L* = 1, 2, 4, and 8) and the coefficient of variation in fixation times with *L* = 8 in the multi−locus simulation of the TP model. Error bars show two times the standard errors. A total of 100 replicates of simulations were run under soft selection with Φ = 2 (cyan; *a*_*S*_ = 0.1, *b*_*S*_ = 0, *a*_*U*_ = 0.25, *b*_*U*_ = *a*_*V*_ = *b*_*V*_ = 0.05), hard selection with Φ = 1 (brown; *a*_*S*_ = 0.1, *b*_*S*_ = 0, *a*_*U*_ = 0.1, *a*_*V*_ = 0, *b*_*U*_ = *b*_*V*_ = 0.1), or hard selection with Φ = 0.75 (orange; *a*_*S*_ = 0.2, *b*_*S*_ = 0, *a*_*U*_ = 0.25, *a*_*V*_ = 0.1, *b*_*U*_ = *b*_*V*_ = 0.05). Other parameters are identical to those in Figure 4A.

With hard selection, however, 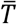 decreased as *L* increased. Furthermore, the coefficient of variation, the standard deviation of *T*_*i*_’s divided by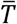, was significantly smaller with hard selection than with soft selection (Fig. 5B). As expected for hard selection, the amplitude of the population size fluctuation progressively increased as the number of loci fixed for *A*_2_ increased (Fig. 4A). These results clearly demonstrate that positive feedback between demographic and selective fluctuations under hard selection led to a great amplification of population size fluctuation; an initial fluctuation in 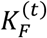 positively selected the mutant allele *A*_2_ at one locus whose fitness fluctuates in positive correlation with 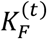, leading to the fixation of *A*_2_ and therefore a larger fluctuation of the field (given by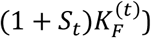, which makes it easier for the *A*_2_ allele at another locus to be positively selected. This process could continue to increase the fluctuation of the field until its carrying capacity is given by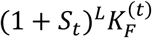, where *L* is the number of loci available in the genome for mutation into *A*_2_. There was also a moderate effect of the recombination rate in accelerating the multi-locus fixation of *A*_2_ with hard selection (Table S2). Fixation times generally increased with decreasing recombination rates for both soft and hard selection, possibly because tighter linkage increased the occurrence of clonal interference.

### Larval-subadult-adult (LSA) model

Althoug the refuge in the TP model is expected to play the role of various demographic/genetic elements that were previously shown to produce the storage effect, such as seed banks, diapausing stages, or a particular life-stage in a population with overlapping generations, the noncovarying storage effect found in the TP model may not be replicated in other population structures as the nature of refuge (i.e. whether it undergoes its own reproduction) and the pattern of ‘migration’ in and out of the refuge can be qualitatively different. I therefore investigate an explict discrete-time model of an age-structured population with overlapping generations. Three stages – ‘larval’, ‘subadult’, and ‘adult’ – in the life cycle of a haploid species are assumed. Time is counted in intervals of the length required for one life stage to advance to the next one. At the end of each interval, the final size of larval population is regulated to its carrying capacity (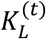at time *t*). Probabilities for an individual to advance from larval to subadult and from subadult to adult stages are given by *e*_*L*_ and *e*_*S*_, respectively. Therefore, the sizes of the subadult and adult populations are expected to be 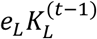and 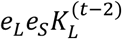at time *t*. 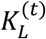is assumed to vary randomly over time with no autocorrelation. Specifically it is modeled to be 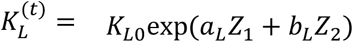, where *Z*_1_ and *Z*_2_ are standard normal variables drawn each time.

The population is made of haploids carrying either the *A*_1_ or *A*_2_ allele at each of *L* linked loci. The (marginal) fitness of *A*_2_ relative to *A*_1_ at locus *j* is given by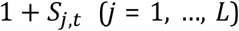, where 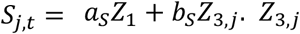. is drawn independently for different loci at each interval *t*. Then, the fitness of haplotype *i*, 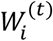, is given by the multiplication of allelic fitness across all loci. It is assumed that this haplotype fitness determine the viability in the larval stage only; the carriers of two alleles are not different either in the probability of advancing to the next stage or in fertility in the subadult and adult stages. If there are only larval and subadult stages, they constitute a population that reproduces in effectively discrete generations. The addition of the adult stage provides a storage for alleles regardless of their reproductive success in the previous (subadult) stage. Therefore, the adult stage plays the role of the refuge and is an example of a noncovarying storage because its size is independent of the concurrent subadult stage.

Multi-locus simulations, allowing not only genetic drift in the reproduction step but also stochastic advances from one to the next life stage (see Appendix B for method), were performed with limited sets of parameters, each starting with a population fixed for haplotype 0, which carries *A*_1_ at all loci. The evolutionary dynamics of the *A*_2_ alleles in each simulation run over 50,000 time units was summarized by the mean heterozygosity and the proportion of time *A*_2_ remained close to relative frequency 0 or 1 (Table S3). Major results produced in the TP model – multi-locus balancing selection under soft selection or positive directional selection on *A*_2_ under hard selection – were readily obtained with this model. In agreement with Kim (2023) and the TP model, the relative strength of demographic vs. fitness fluctuation determined the outcome; when we define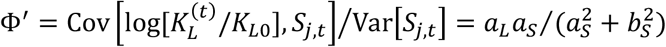, which is analogous to Φ in the TP model, positive selection on rare *A*_2_ occurred when Φ’ was above 0.7. With soft selection and above this threshold of Φ’, the frequency of *A*_2_ underwent oscillations that are indicative of balancing selection (Fig. S3). Then, as Φ’ continued to increase, balancing selection transitioned into positive directional selection. With hard selection, positive directional selection occurred once Φ’ increased above the threshold. Exemplary replicates show that, as the *A*_2_ alleles reach fixation across loci, fluctuations in the sizes of all three life stages become amplified (Fig. 4B, Fig. S3). Negative (positive) correlation between mean fixation times and the number of loci under hard (soft) selection and the clustering of fixation times with hard selection were also oberved in other sets of simulations (Fig. S4), which basically replicated the results from the TP model. This confirmed that positive feedback between demographic and fitness fluctuations under hard selection occurs in both models. In conclusion, all major aspects of the noncovarying storage effect found in the TP model were shown to arise in the LSA model as well.

## DISCUSSION

A key assumption in this study is that a tradeoff (antagonistic environmental pleiotropy; Zhang 2023) commonly occurs for the fitness effects of a mutant allele between different time points in the environmental fluctuation. Namely, a mutant phenotype that increases the reproductive output relative to others when the environment is generally favorable is likely to decrease it relative to others during an unfavorable period. Such a mutant allele increases the variance of the offspring number over time while the mean may remain unchanged. For example, as postulated by Slatkin (1974), a mutation that makes a female bird lay 10 eggs in one batch instead of spreading them over time in a randomly and rapidly fluctuating environment will increase the variance of her fitness while the mean remains the same. Setting the fitness of a mutant to 1 + *S*_*t*_ with E[*S*_*t*_] = 0 models such a tradeoff. Classical theory predicts that such a mutant (termed an arithmetic-mean-preserving or AMF allele in Kim (2023)) is eliminated from a population as its geometric mean fitness is less than 1.

Kim (2023) and this study however found that the AMF mutant at a low frequency is positively selected (with both soft and hard selection) if a correlation between demographic and fitness fluctuations exists in the presence of a refuge whose carrying capacity oscillates in a narrower range and/or out of phase relative to the field. A heuristic argument for this noncovarying storage effect on the fitness-amplifying mutant allele may be given. In the TP model, the mean number of offspring per parent, *i*.*e*. the absolute fitness, of the rare mutant in the field is given by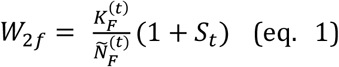, where 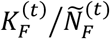 is the absolute fitness given under demographic fluctuation alone. With positive correlation between 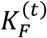and *S*_*t*_, the fluctuation of *W*_2*f*_ becomes greater than that of 1 + *S*_*t*_ and, more importantly, its arithmetic mean can now exceed 1. Still, the geometric mean of 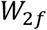 is not greater than 1 due to its fluctuation over time. However, the presence of a refuge can reduce the variance of mutant’s fitness averaged across the whole population such that the geometric mean of this average fitness becomes greater than 1. Here, to generate this dampening effect, the refuge should not undergo the same fluctuation in size with the field; the absolute fitness in the refuge 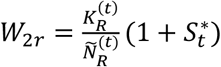in eq. (1) should fluctuate differently than 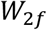. With parameter values used in of the TP model, 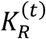was given to covary incompletely with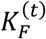. In the LSA model, the fluctuation of the refuge (the adult stage) is uncorrelated to the fluctuation of the field (the subadult stage). In Kim (2023), the seed bank fluctuates in size similar to the subadult and adult stages in the LSA model. In summary, such refuges, or noncovarying storages, turn the heightened arithmetic mean of the mutant allele raised by the demographic fluctuation of the field into an elevation of the geometric mean in the total population.

Not all refuges in the previously proposed mechanisms of the storage effect are noncovarying. For example, it was suggested that if a gene is expressed in one sex in a dioecious species the other sex could be a refuge (Reinhold 2000; Kim 2023). However, as equal numbers of male and female gametes are used in reproduction, the subpopulation sizes of reproductive males and females are effectively equal while the sex ratio of the adults may fluctuate. Therefore, as the size of the field relative to the refuge does not fluctuate at all, the noncovarying storage effect should not arise for the genes of the sex-limited expressions.

One may also model the fluctuating fitness of a mutant by exp[*S*_*t*_], making the mutant (termed a geometric-mean-preserving or GMF allele) quasi-neutral as its geometric mean fitness is identical to that of the wild-type allele even without demographic fluctuation. Analyzing the model of cyclic environmental oscillation, Kim (2023) showed that the parameter range of positive selection for a GMF allele is wider than that for an AMF allele. Similarly, GMF alleles should be subject to the noncovarying storage effect under broader conditions. However, a GMF allele may be considered simply a beneficial allele as the arithmetic mean of offspring number is greater than the wild-type allele. Such a mutation is probably rare and therefore was not considered in this study. One should however note that GMF alleles might be still important for genetic variation in nature since they can be maintained in balanced polymorphism in the conventional models of the storage effect, for example without demographic fluctuations or in the case of sex-limited gene expressions.

Analytic approximation for the one-locus TP model predicts that, under soft selection or strict density regulation that keeps the population size to the carrying capacity of the field at a given time regardless of its genetic composition, balancing selection arises if the demographic fluctuation (a major contributor to Cov[*S*, log[*K*_*F*_/*K*_*R*_]]) is not so weak or strong compared to fitness fluctuation (Var[*S*] in case of *S*^∗^ = 0). Here, balancing selection is defined as a force that generates polymorphism at a level greater than expected for neutral alleles. Multi-locus simulation showed that balanced polymorphism can be simultaneously maintained at many loci, as expected since the demographic fluctuation (together with the presence of refuge) that turns the geometric mean fitness of an AMF allele from < 1 to > 1 should apply equally to all loci in the genome. Previously, Park and Kim (2019) showed that, in the presence of a refuge but without fluctuation in population size, polymorphism can arise at multiple loci that contribute to the expression of a phenotype under fluctuating selection. However, the number of loci where balanced polymorphism emerged increased very slowly as the total number of loci in the model increased. In contrast, in the current simulation, balanced polymorphism was observed at all loci available for mutations to arise. However, the model containing only up to 10 loci was examined, due to the limitation of the simulation method used, and the 10-loci polymorphism at each locus was generally lower than the level observed in the one-locus simulation. This is probably because the segregation of alleles at multiple loci elevates variance in the offspring number well above that of the Poisson distribution, thus increasing genetic drift that weakens the positive selection on rare alleles. The possibility that this mechanism can maintain the genome-wide polymorphism over 1,000 loci, currently observed in several natural populations (Bergland, et al. 2014; Johnson, et al. 2023), should be addressed by a simulation on a large scale in terms of both the number of loci and the census population size.

This study focused more on the other evolutionary outcome of the noncovarying storage effect, the fixation of mutations that amplify fitness fluctuations. In the TP model, the condition for the fixation is less restrictive compared to that for balanced polymorphism, because it requires relatively weaker selection (smaller variance in relative fitness) for a given demographic fluctuation to satisfy, assuming *S*^∗^ = 0, Φ > 1 + 1/(2*ρ*) with soft selection and Φ > 1/(2*ρ*) with hard selection. Similar ranges of conditions (for Φ’) seem to apply for the LSA model (Table S3). Demographic fluctuation that generates the noncovarying storage effect should apply equally to all loci in the genome. Therefore, fixations occur at all available loci on which proper mutations (*i*.*e*. the *A*_2_ allele) arise, as observed in the multi-locus simulation. With soft selection, the pattern of demographic fluctuation that generates the noncovarying storage effect does not change over time regardless of any evolutionary change at any locus. However, with hard selection, fluctuation in the relative size of the field generating the nocovarying storage effect at a given locus intensifies as more fluctuation-amplifying mutations reach fixation at other loci. Since the carrying capacity of the field in the TP model is effectively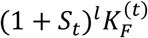, where *l* is the number of loci where *A*_2_ reached fixation, and wider fluctuation generates stronger positive selection on the rare *A*_2_ (larger E[Δ_2_] in Appendix A), fixation at one locus facilitates the subsequent fixations at other loci. This positive feedback between demographic fluctuation and positive selection drastically increases the amplitude of fluctuation in the total population size (Fig. 4). How much the amplitude will increase should depend on the number of loci at which mutations subject to hard selection arise. There should be multiple independent ways in which changes in mutants’ relative fitness can be translated into changes in the mean absolute fitness of the entire population. For various life history traits, one may imagine phenotypes (smaller eggs, smaller adult body, earlier onset of first reproduction, etc.) that increase the absolute number of surviving offspring during a favorable, resource-rich period but decrease the number in a bad period while not affecting the amount of resources that the carriers of the alternative, ancestral alleles acquire. Then, many loci may be available for hard selection, the combined effect of which may lead to a very large fluctuation in population size.

The size of a population and its fluctuation should be governed not only by extrinsic factors (the environmental variables such as temperature and parasite prevalence) but also by intrinsic factors (species-specific responses to those extrinsic factors). The intrinsic factors such as various life-history traits are the product of the species’ evolutionary history. For example, phenotypic plasticity responding to environmental cues can reduce the amplitude of the population size fluctuation, thus acting as a mechanism of population regulation (Caswell 1983; Reed *et al*. 2010), which according to the classical theory is adaptive as it enhances long-term reproductive success by increasing the geometric mean fitness (Gillespie 1977; Pfister 1998). If life history traits evolved in the direction toward suppressing random or cyclic fluctuations in the population size, fluctuations observed in nature should be interpreted as the effect of variable environments that the evolved mechanisms of population regulation were unable to contain. However, this study suggests that evolution can proceed in the opposite direction. A large random fluctuation in population size can be the product of adaptive evolution driven by positive feedback between demographic and selective fluctuations. When the degrees of population size fluctuation are different across species, the eco-evolutionary dynamics proposed in this study might be particularly important for those exhibiting large fluctuations. Interestingly, annual desert plants experiencing wider fitness fluctuations after germination have greater proportions of their population remaining in the seed bank (Gremer & Venable 2014). This observation is in agreement with the analytic results in this study because a larger seed bank corresponds to a larger value of ρ, which makes it easier for the fixation of the fluctuation-amplifying alleles (i.e., lowering the threshold of Φ for positive selection).

## DATA AVAILABILITY

Mathematica notebook files for conducting all simulations reported are available from https://github.com/YuseobKimLab/RandomFluctuationEcoEvo

## ACKNOWLEDGMENT

This study was supported by the National Research Foundation (NRF) grant 2020R1A2C1009261 funded by the Korean government.

## Appendix A Derivation of conditions for balancing and directional selection on *A*_2_ in the TP model

Here only deterministic changes in allele frequencies are tracked as a very large population is assumed. At the end of reproduction but before migration into subpopulations at generation *t* - 1, there are *n*_1_ and *n*_2_ copies (individuals) of *A*_1_ and *A*_2_ in the entire population. After migration, the numbers of *A*_1_ and *A*_2_ copies are *n*_1*F*_ = (1 − *r*)*n*_1_ and *n*_2*F*_ = (1 − *r*)*n*_2_ in the field and *n*_1*R*_ = *rn*_1_ and *n*_2*R*_ = *rn*_2_ in the refuge. Therefore, the relative frequencies of alleles are identical between the field and refuge before reproduction at the beginning of generation *t*.

Soft selection is first considered. Then, each *A*_2_ allele produces 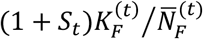 copies of descendants, where 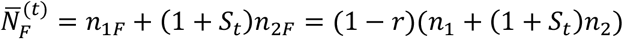. Since *n*_1_ » *n*_2_ after *A*_2_ appears by mutation, 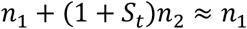 and 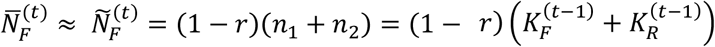. Similar approximations apply to the refuge. Let *n’*_2_ be the expected number of *A*_2_ in generation *t*. This leads to eq. (1). Namely,

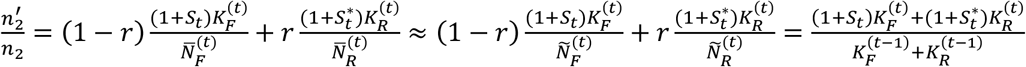

Then, with 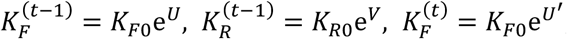, and 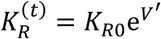

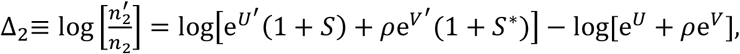

where *S* = *S*_*t*_, 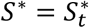, and *ρ* = *K*_*R*0_/*K*_*F*0_. If E[Δ_2_] > 0, where the expectation is over demographic and fitness fluctuations, the rare allele *A*_2_ is expected to increase its copy number over time. As the expectations of *U, V, U’, V’, S*, and *S*^*\**^ are all zero and non-zero correlations exist only among *U’, V’, S*, and *S*^*\**^ and between *U* and *V*, from the Taylor expansion of Δ_2_ up to the second order, it can be shown that

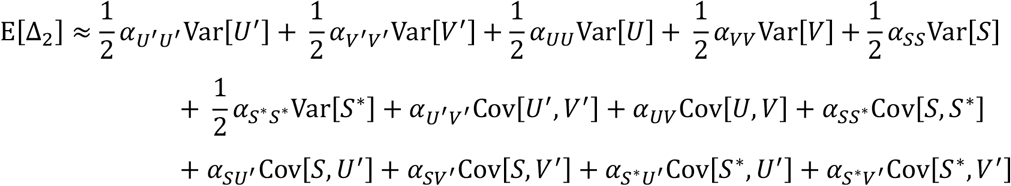

where 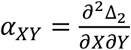 is the Taylor coefficient for variables *X* and *Y* evaluated at (*U, V, U*^′^, V^′^, S, *S*^∗^) = (0,0,0,0,0,0). Since 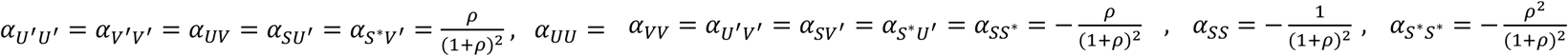, Var[*U*] = Var[*U*^′^], Var[*V*] = Var[*V*^′^], and Cov[*U, V*] = Cov[*U*^′^, V^′^], the above equation is simplified to

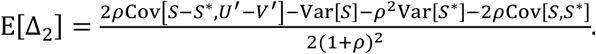

Therefore, the condition for the *A*_2_ allele being positively selected (i.e. invading the population initially fixed for *A*_1_), is E[Δ_2_] > 0 or

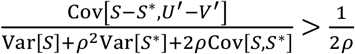

assuming (Var[*S*] + *ρ*^2^Var[*S*^∗^])/*ρ* > −2Cov[*S, S*^∗^]. If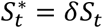, the above condition becomes

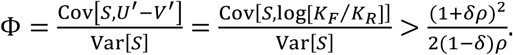

A similar result can be obtained for the condition for the *A*_1_ allele to invade a population initially fixed for *A*_2_. The expected change in the number of *A*_1_, *n’*_1_/*n*_1_, is obtained with a procedure identical to that leading to eq. (1) except that the relative fitness is flipped. Therefore

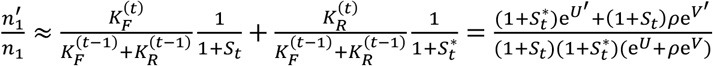

for *n*_1_ ≪ n_2_, the dynamics of the rare *A*_1_ allele depends on the expectation of

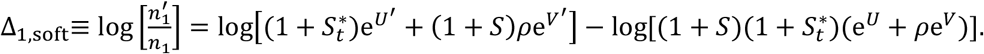

Applying the quadratic approximation used for obtain E[Δ_2_] above, one obtains

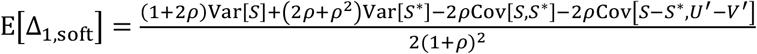

The condition for the rare *A*_1_ allele to increase in frequency is [EΔ_1,soft_]> 0 or

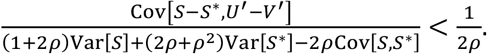

If 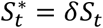, the above condition becomes

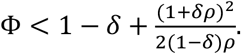

With hard selection, the frequency change of *A*_2_ when rare is again given by eq. (1). Therefore, the condition for *A*_2_ invading the population fixed for *A*_1_ (E[Δ_2_] > 0 above) is no different from that for soft selection. However, because the carrying capacities of the field and refuge fixed for *A*_2_ at generation *t* is given by 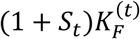and 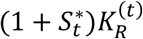and 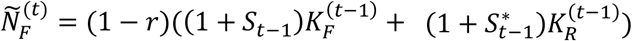 and 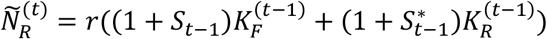under hard selection, the opposite condition for *A*_1_ invading *A*_2_ is given by

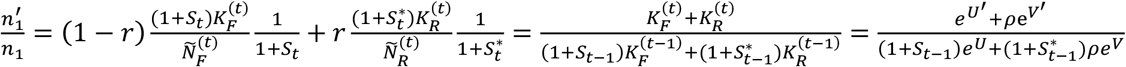

Then,

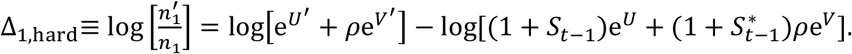

Now, given that 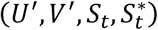and 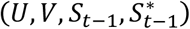are independent samples of a joint distribution, the expectation of −Δ_1,hard_ is exactly that of Δ_2_ above. Therefore, the condition for the fixation of *A*_2_ (E[Δ_1,hard_]< 0) is exactly the condition for positive selection on a rare *A*_2_ (E[Δ_2_] > 0).

## Appendix B Methods of stochastic simulation

Stochastic simulations of the single- and multi-locus eco-evolutionary models were implemented in Mathematica ver. 11.3. In both TP and LSA models, an individual has *L* loci that are arranged linearly with probability *c* for recombination between adjacent loci per generation. The simulation keeps track of two sets of lists: one containing the sets of haplotypes carried by non-zero individuals in the field and in the refuge (***H***_F_ and ***H***_R_) and the other containing the corresponding numbers of individiuals (***N***_F_ and ***N***_R_).

In the TP model, generation *t* begins after the migration step (re-distribution of individuals produced in the previous generation into the field and refuge) is completed. Haplotype numbers are first updated by mutation and recombination. A Poisson variate with mean *NL*μ, where *N* is the current size of a subpopulation at hand, is drawn as the total number of mutation events and each event is assigned to haplotypes and loci proportional to their current frequencies. Similarly, for *L* > 1, a Poisson variate with mean *N*(*L*-1)*c* is drawn as the total number of recombination events in the subpopulation. For each recombination event, two haplotypes are chosen proportional to their frequencies and one of the *L*-1 intervals is chosen to be the position of crossing-over between the two haplotypes. Newly created or lost haplotypes during mutation and recombination steps are tracked by updating (***H***_F_, ***H***_R_) and (***N***_F_, ***N***_R_). Then, the updated lists are further updated in the step of reproduction according to their fitness. Under soft selection, the number of individuals with haplotype *i* born in the next generation is Poisson distributed with mean 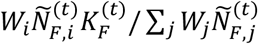in the field and 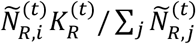 in the refuge, where *W*_*i*_ is the relative fitness of haplotype *i* and 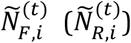 is the number of haplotype *i* individuals in the field (refuge) after the haplotypes are updated by mutation and recombination. Under hard selection, the corresponding mean number in the field is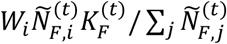. Finally, when there are *N*_F_ and *N*_R_ individuals in the field and refuge, Poisson variates with the mean *N*_F_*m*_FR_ (*N*_R_*m*_RF_) truncated at the upper bound *N*_F_ (*N*_R_) is drawn to determine the numbers of migrants from field to refuge (from refuge to field). Migrants are chosen proportional to the haplotype frequencies and then merged with the residents of the destination. This completes a single-generation iteration in updating haplotype lists.

Simulation for the LSA model is similarly performed by tracking the haplotypes. Let *l*_*i*_, *m*_*i*_, and *n*_*i*_ be the number of individuals with haplotype *i* (*i* = 1, …, 2^*L*^) at the larval, subadult, and adult stages at time *t*. The following steps are taken to produce the corresponding numbers, *l*^*i*^’, *m*^*i*^’, and *n*^*i*^’, at time *t*+1. As in the TP model, bidirectional mutations with probability μ per locus and recombination with probability *c* for adjacent loci occur but in subadult and adult stages only. Recombination must occur among subadults and among adults only. The updated numbers of individuals by haplotypes in subadults and adults after mutations and recombination are *m*_*i*_^*^, and *n*_*i*_^*^. Next, the numbers of individuals by haplotypes in the next generation are obtained. *l*_*i*_’ is given by a Poisson number with mean 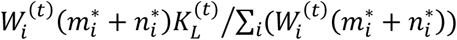under soft selection and 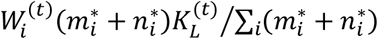under hard selection. *m*_*i*_’ and *n*_*i*_’ are given by Poisson numbers with mean *e*_*L*_*l*_*i*_ and *e*_*L*_*e*_*S*_*m*_*i*_^*^, respectively.

**Table S1.**
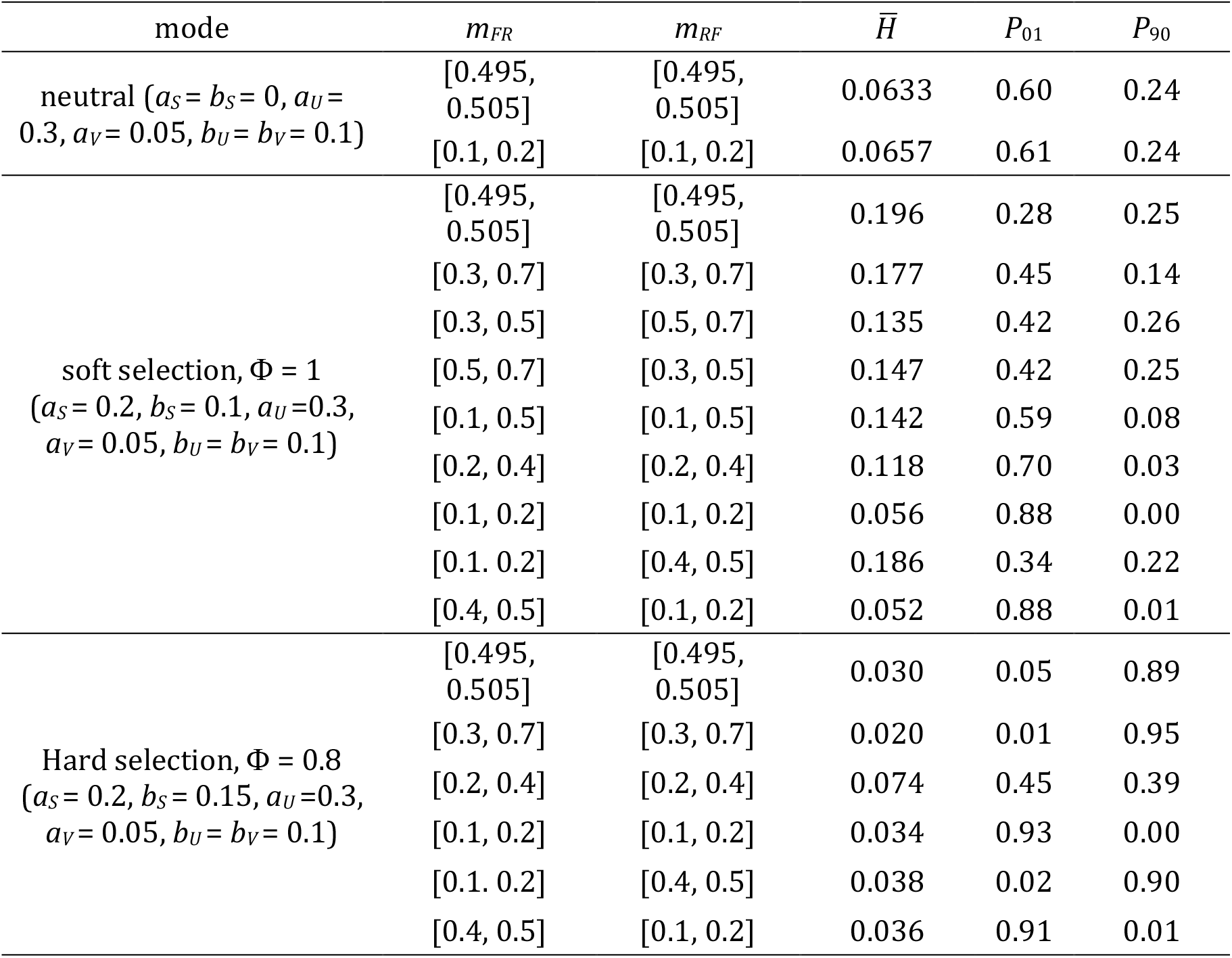
One-locus simulation with variable migration rates in the TP model.

**Table S2.**
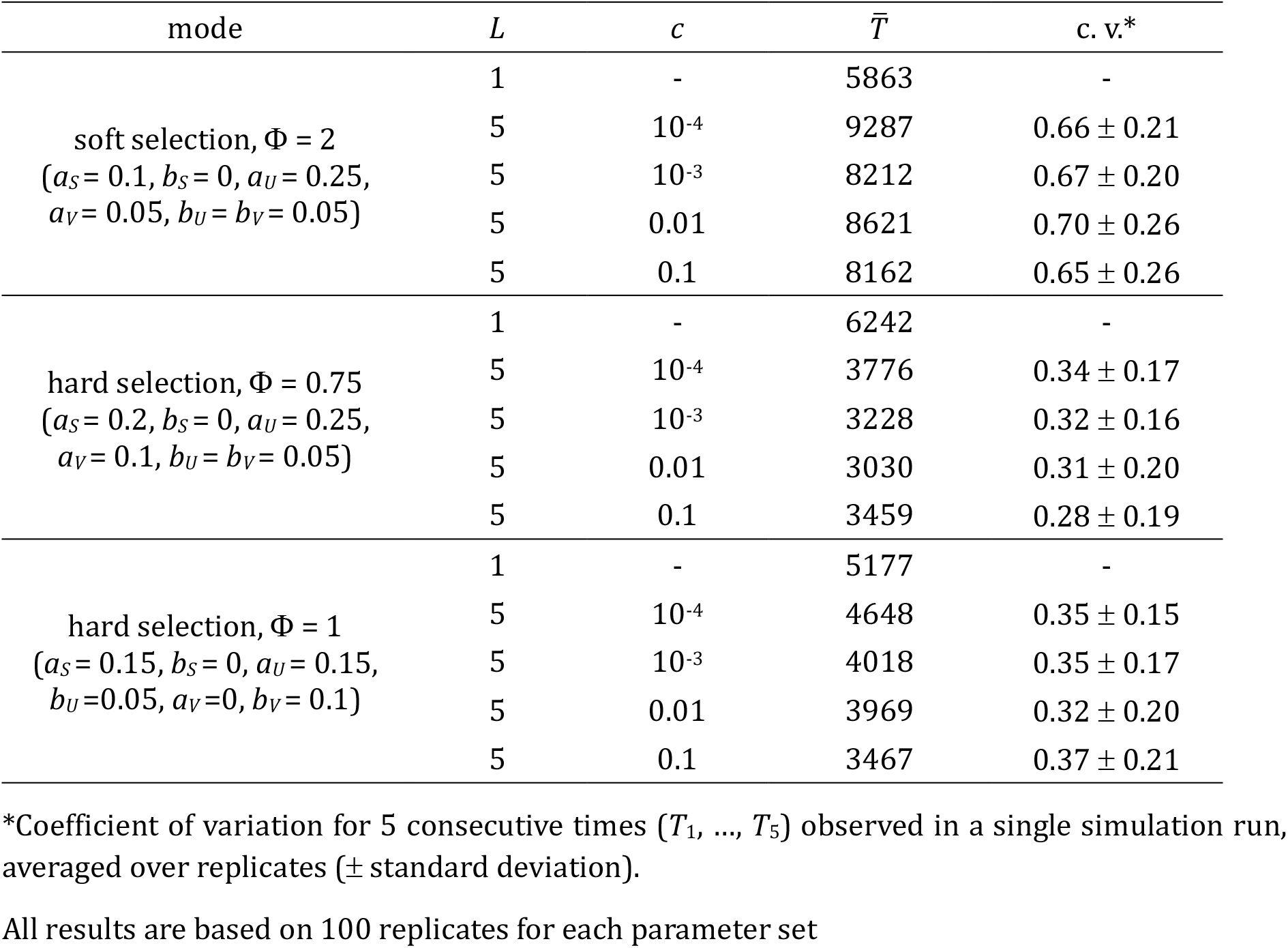
Mean time to the fixation (*q* > 0.9) of *A*_2_ in the multi-locus simulation of the TP model.

**Table S3.** Long-term polymorphism in multi-locus simulations of the LSA model.

| mode | $a_s$ | $b_s$ | $\Phi'$ | $\bar{H}$ | $P_{01}$ | $P_{90}$ |
| --- | --- | --- | --- | --- | --- | --- |
| neutral | 0 | 0 | - | 0.0757 | 0.42 | 0.40 |
| soft<br>selection | 0.05 | 0.2 | 0.294 | 0.013 | 1 | 0 |
|  | 0.1 | 0.2 | 0.5 | 0.017 | 0.96 | 0 |
|  | 0.1 | 0.15 | 0.769 | 0.040 | 0.91 | 0.00 |
|  | 0.2 | 0.1 | 1 | 0.152 | 0.63 | 0.08 |
|  | 0.2 | 0 | 1.25 | 0.116 | 0.31 | 0.41 |
|  | 0.15 | 0.05 | 1.5 | 0.054 | 0.17 | 0.71 |
|  | 0.1 | 0.05 | 2 | 0.023 | 0.22 | 0.73 |
| hard<br>selection | 0.05 | 0.2 | 0.294 | 0.006 | 0.99 | 0 |
|  | 0.1 | 0.2 | 0.5 | 0.013 | 0.97 | 0 |
|  | 0.1 | 0.15 | 0.769 | 0.009 | 0.22 | 0.76 |
|  | 0.2 | 0.1 | 1 | 0.003 | 0.07 | 0.93 |
|  | 0.15 | 0.087 | 1.25 | 0.003 | 0.24 | 0.76 |
|  | 0.15 | 0.05 | 1.5 | 0.005 | 0.06 | 0.92 |

- Each simulation ran for 50,000 time units, except for the neutral simulation that ran 500,000 time units.
- Other parameters: $\mu = 2 \times 10^{-5}$ , $K_{L0} = 5000$ , $a_L = 0.25$ , $b_L = 0.1$ , $e_L = 0.7$ , $e_S = 0.7$ , $L = 4$ , $c = 0.03$ .

**Figure S1.**
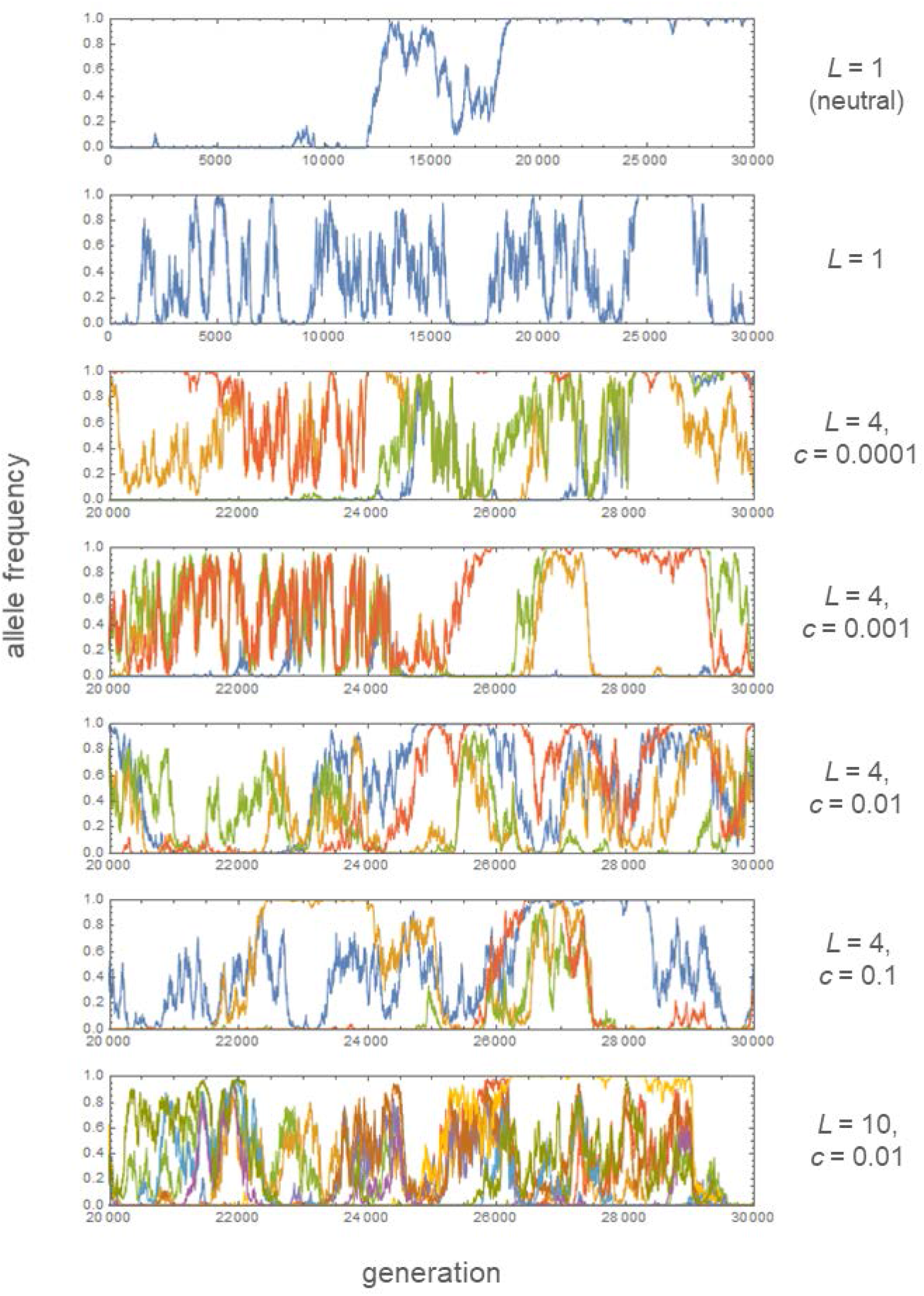
Exemplary trajectories of allele (*A*_2_) frequencies in simulations. Allele frequencies are plotted in different colors for different loci. The number of loci (*L*) and recombination rate (*c*) are shown on the right side of graphs. Other parameters: *K*_*R*0_ = *K*_*F*0_ = 1000, μ = 2×10^−5^, Φ = 1 (*a*_*S*_ = 0.2 [0 for neutral], *b*_*S*_ = 0.1, *a*_*U*_ = 0.3, *b*_*U*_ = 0.1, *a*_*V*_ = 0.05, *b*_*V*_ = 0.1).

**Figure S2.**
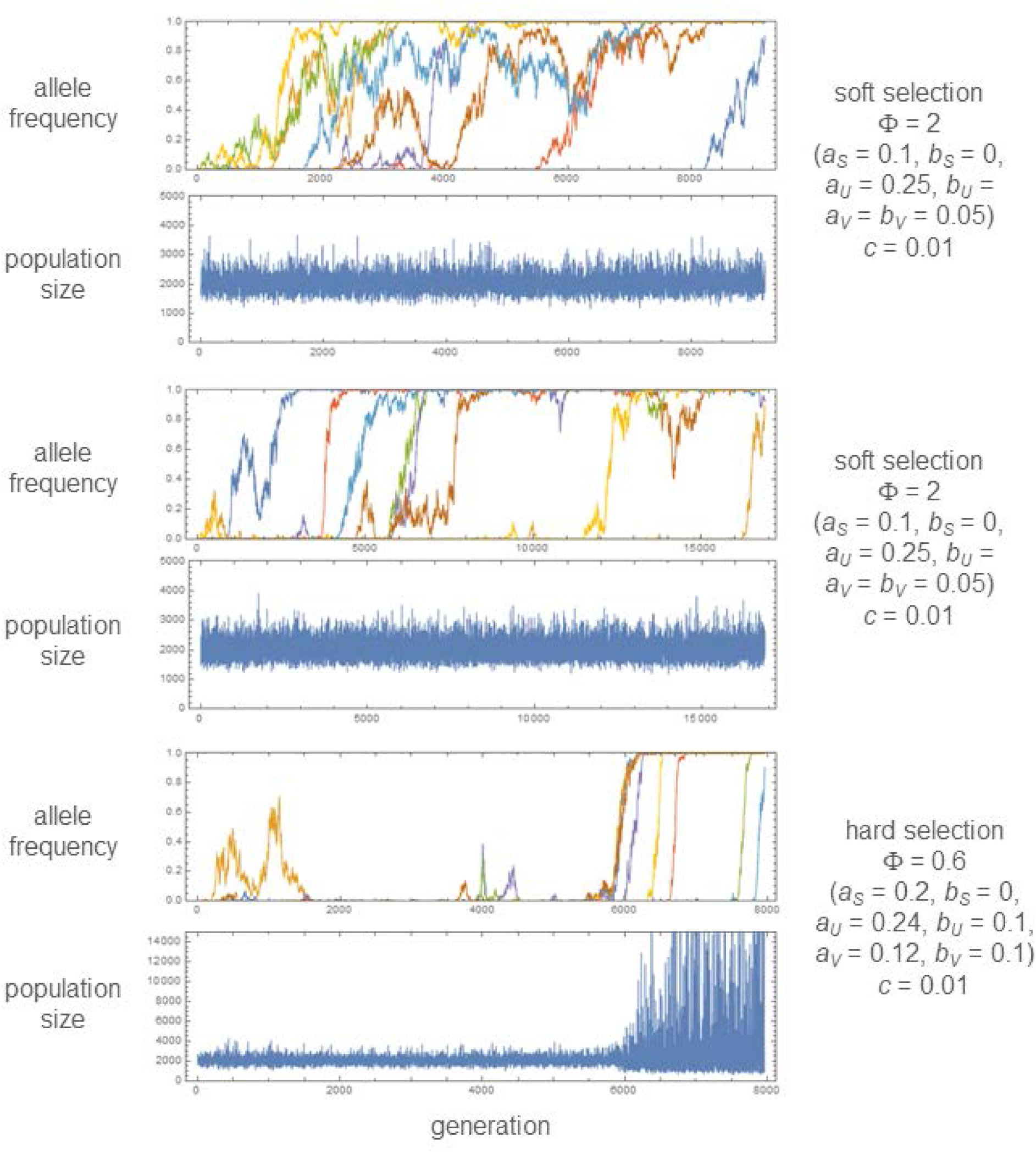
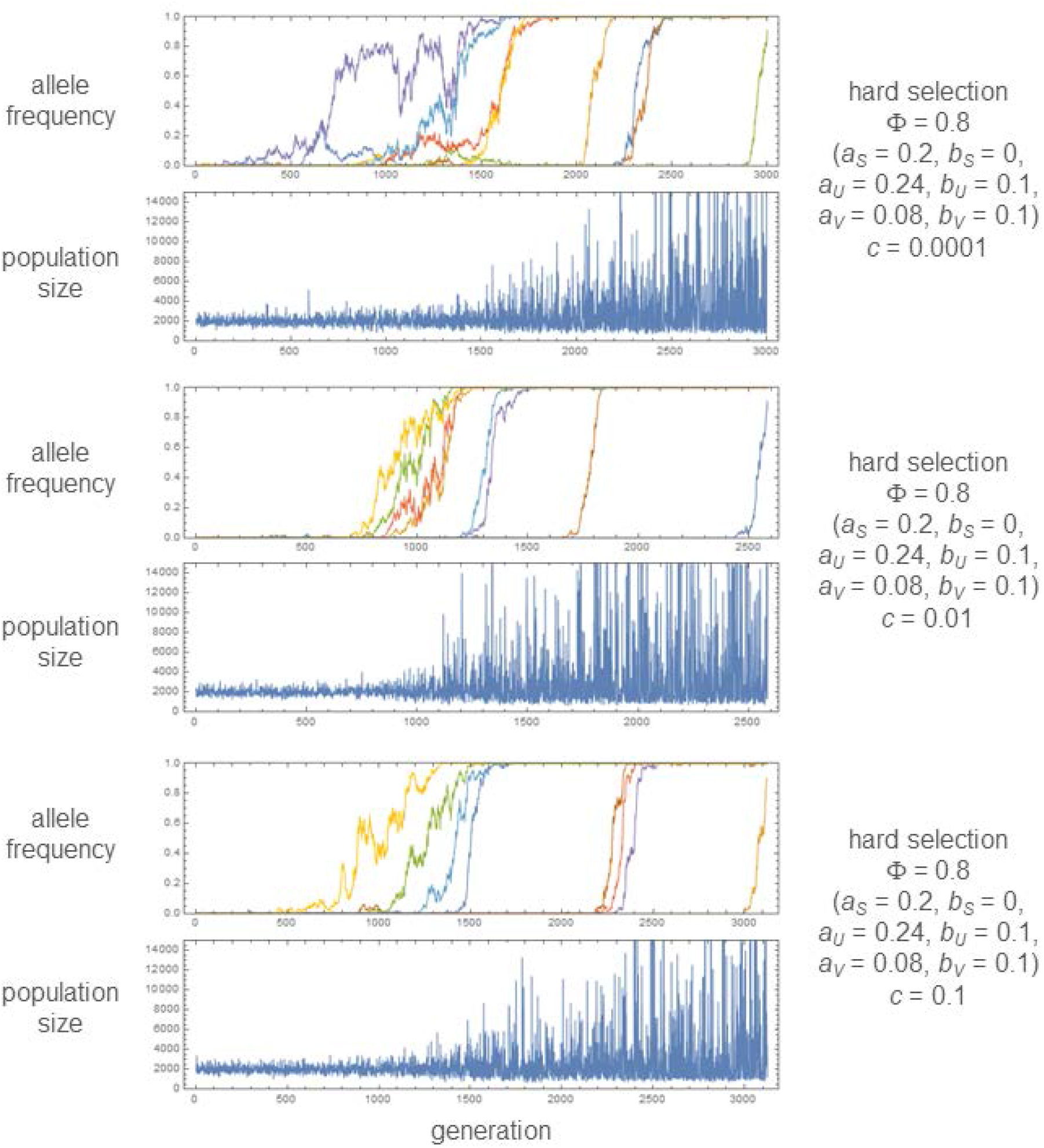
Examples of simulation runs in which the mutant allele (*A*_2_) reaches fixation sequentially at all 8 loci used in the TP model. The change of population size is shown below that of allele frequencies, plotted in different colors for different loci. Parameter values are shown on the right side of graphs. Other parameters: *K*_*R*0_ = *K*_*F*0_ = 1000, μ = 2×10^−5^.

**Figure S3.**
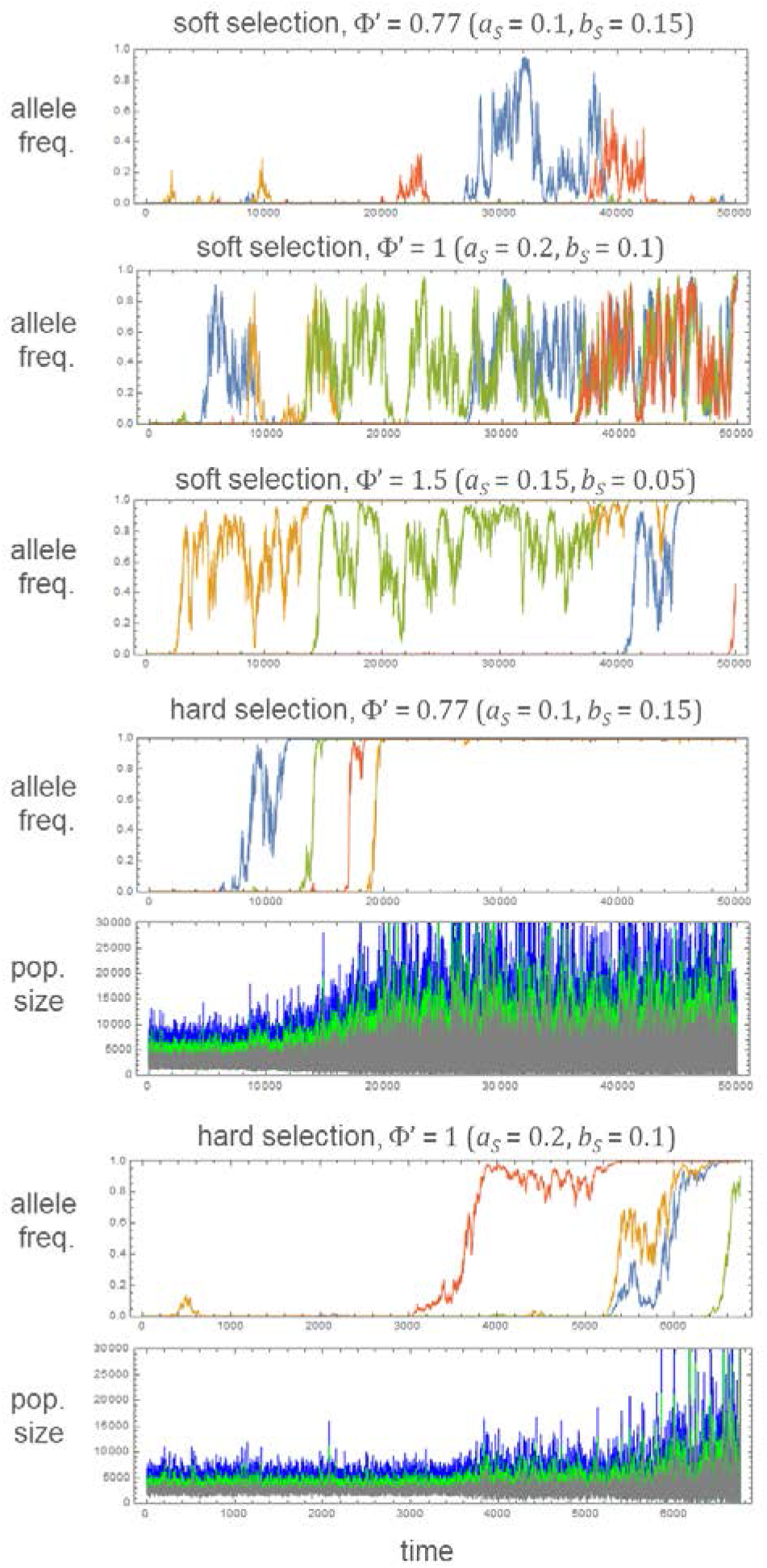
Exemplary allele frequency changes in the simulation of the LSA model. Each simulation started with a population fixed for the *A*_1_ allele and ran for 50,000 time units, except for the case of hard selection with Φ’ = 1. The *A*_2_ allele appears by recurrent mutations at each of *L* = 4 loci. Allele frequencies are plotted in different colors for different loci. For hard selection, the change of population size (blue: larval, green: subadult, gray: adult) is shown below the allele frequency trajectories. Other parameters: *K*_*L*0_ = 5000, *a*_*L*_ = 0.25, *b*_*L*_ = 0.1, *e*_*L*_ = *e*_*S*_ = 0.7, μ = 2×10^−5^, *c* = 0.03.

**Figure S4.**
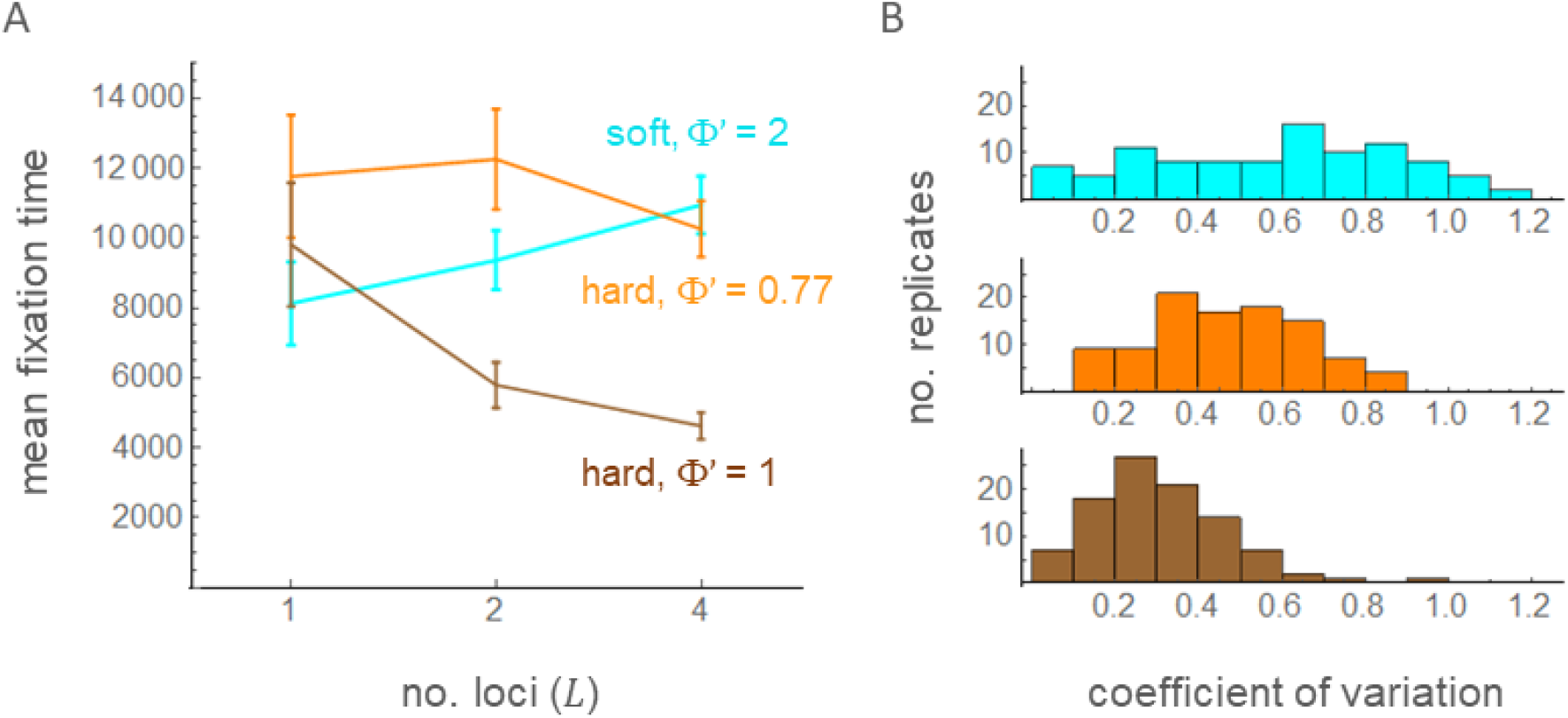
Mean fixation time 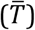 with varying number of loci (A; *L* = 1, 2, and 4) and the coefficient of variation in fixation times with *L* = 4 in the multi-locus simulation of the LSA model. Error bars show two times the standard errors. A total of 100 replicates of simulations were run under soft selection with Φ’ = 2 (cyan; *a*_*S*_ = 0.1, *b*_*S*_ = 0.05, *a*_*L*_ = 0.25, *b*_*L*_ = 0.1, μ = 3×10^−5^), hard selection with Φ’ = 1 (brown; *a*_*S*_ = 0.2, *b*_*S*_ = 0.1, *a*_*L*_ = 0.25, *b*_*L*_ = 0.1, μ = 2×10^−5^), or hard selection with Φ’ = 0.77 (orange; *a*_*S*_ = 0.1, *b*_*S*_ = 0.15, *a*_*L*_ = 0.25, *b*_*L*_ = 0.1, μ = 3×10^−5^). Other parameters are identical to those in Figure S3.

